# Dynamical modeling reveals RNA decay mediates the effect of matrix stiffness on aged muscle stem cell fate

**DOI:** 10.1101/2023.02.24.529950

**Authors:** Zachary R. Hettinger, Sophia Hu, Hikaru Mamiya, Amrita Sahu, Hirotaka Iijima, Kai Wang, Gabrielle Gilmer, Amanda Miller, Gabriele Nasello, Antonio D’Amore, David A. Vorp, Thomas A. Rando, Jianhua Xing, Fabrisia Ambrosio

**Author notes:** Corresponding Author: Fabrisia Ambrosio, PhD, MPT, Suite 5.303, MGH CNY 149. 149 13^th^ St, Charlestown, MA 02129. Institute for advanced research, Nagoya University, Nagoya, Japan. Biomedical and Health Informatics Unit, Graduate School of Medicine, Nagoya University, Nagoya, Japan.

## Abstract

Loss of muscle stem cell (MuSC) self-renewal with aging reflects a combination of influences from the intracellular (e.g., post-transcriptional modifications) and extracellular (e.g., matrix stiffness) environment. Whereas conventional single cell analyses have revealed valuable insights into factors contributing to impaired self-renewal with age, most are limited by static measurements that fail to capture nonlinear dynamics. Using bioengineered matrices mimicking the stiffness of young and old muscle, we showed that while young MuSCs were unaffected by aged matrices, old MuSCs were phenotypically rejuvenated by young matrices. Dynamical modeling of RNA velocity vector fields *in silico* revealed that soft matrices promoted a self-renewing state in old MuSCs by attenuating RNA decay. Vector field perturbations demonstrated that the effects of matrix stiffness on MuSC self-renewal could be circumvented by fine-tuning the expression of the RNA decay machinery. These results demonstrate that post-transcriptional dynamics dictate the negative effect of aged matrices on MuSC self-renewal.

**Graphical Abstract:** Graphical abstract description: The balance of self-renewal and differentiation in young muscle stem cells (MuSCs) is robust to perturbations of the biophysical microenvironment. In contrast, aged MuSCs are highly sensitive to extrinsic perturbations, and exposure to a youthful microenvironment rejuvenates the self-renewing potential of aged MuSCs by modulating post-transcriptional RNA dynamics.

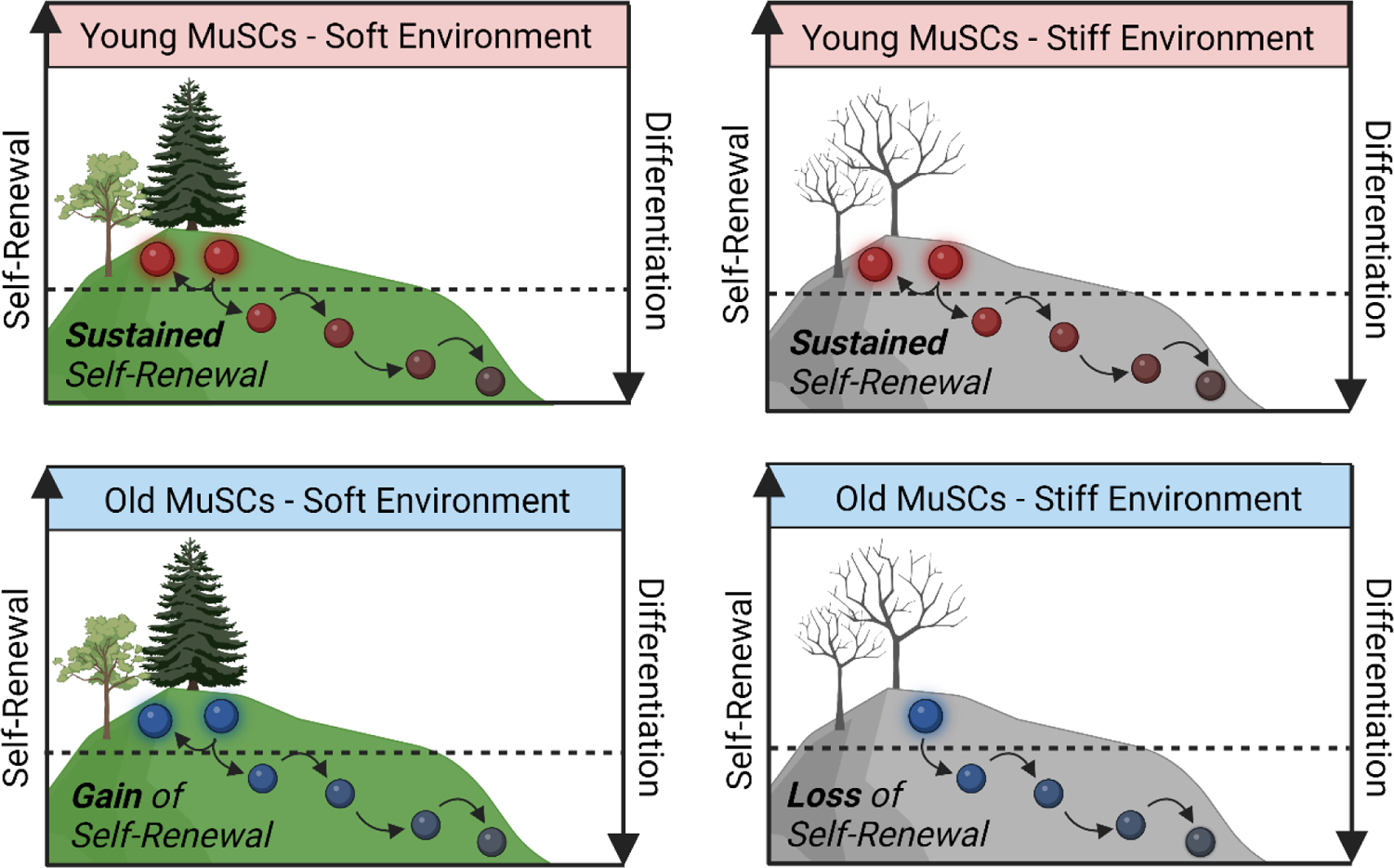

## Introduction

Skeletal muscle regenerative capacity steadily declines with age due to a compromised ability of muscle stem cells (MuSCs) to repair injured muscle and replenish their niche (1, 2). Whereas aberrant lineage specification of aging stem cells has typically been assessed using static measures such as transcript or protein levels, dynamical cell state properties suggests that fate decisions may represent a transition between distinct “attractors”, as has been analogized by Waddington’s landscape (3, 4). Inspired by dynamical systems theory, Waddington depicts the path taken by a marble (i.e., cell) as it rolls down a hill (i.e., the cellular landscape) to represent cellular progression from an undifferentiated to a differentiated state (4). Bifurcations along the path of the marble represent alternative fates. Extending the metaphor, a less common scenario in which the cell starts the path to differentiation but then changes direction to roll up the hill represents a reversion to an undifferentiated state, or self-renewal. Associated with, and perhaps contributing to, Waddington’s epigenetic landscape is the physical landscape of the stem cell niche. Here we asked, *how does the MuSC physical landscape change over time and how do age-related changes to the MuSC landscape affect cellular fate transitions?*

One prominent aspect of the MuSC physical landscape is the extracellular matrix (ECM). ECM mechanical properties represent well-established drivers of stem cell fate, as evidenced by a series of elegant *in vitro* studies demonstrating regulation of stem cell lineage progression via extrinsic mechanical signals (5, 6). For instance, mesenchymal stem cells spontaneously differentiated toward a myogenic lineage when seeded on a soft substrate engineered to mimic the compliance of young, healthy muscle (elastic modulus €: 8 – 17 kPa) (5). On the other hand, osteogenic differentiation was observed when cells were seeded on a stiffer substrate (€: 25 – 40 kPa) (5). Given biophysical characteristics of the ECM are altered with aging (7–9), these *in vitro* findings underscore the potential impact of age-associated changes to the ECM on MuSC lineage specification *in vivo*.

Aged muscle is typically characterized by abnormal ECM deposition and remodeling, contributing to increased tissue stiffness (10–14). Indeed, recent studies from our laboratory have shown decreased collagen tortuosity and a concomitant increase in muscle rigidity along the axis of collagen fibrils in aged muscle (8). These age-related ECM alterations have direct effects on MuSC lineage specification, as we have previously shown MuSCs seeded onto decellularized ECM from aged muscle display an increased fibrogenic conversion compared to cells placed on ECM derived from young muscle (8). However, the cell state regulatory mechanisms that underlie age- and stiffness-dependent MuSC fate decisions are poorly understood. In this study, we tested the hypothesis that age-related alterations in stiffness of the skeletal muscle ECM compromise MuSC lineage progression. Specifically, we employed an array of single cell approaches at the morphological, protein, and RNA levels to determine whether the biophysical properties of the aged muscle physical landscape predispose MuSCs towards differentiation, to test whether the properties of a young niche restore self-renewal in old MuSCs, and to interrogate mechanisms by which MuSCs adopt aberrant fate kinetics upon exposure to an aged niche.

## Results

### Age-related matrix stiffness is associated with aberrant MuSC nuclear morphologies

While numerous reports have highlighted profound age-related biochemical changes to the skeletal muscle ECM (8, 15, 16), comprehensive biophysical characterizations are fewer in number (17). Quantification of the micromechanical properties of muscle is challenging due to the unique ECM architecture and a lack of standardized methodology (18). Moreover, uniaxial testing is limited because it does not capture the out-of-plane properties of muscle (18). To overcome these limitations, we modeled biaxial microscale muscle stiffness using macroscale measurements collected from young (4-6 months) and old (22-24 months) male mice (**Fig. 1A**). The fibril network architectural features of young and old muscle were used to create artificial fiber networks that were stochastically equivalent to experimental samples. On the artificial network, we applied a structural deterministic approach to derive the single fiber Green’s strain, as previously described (18). From these data, we performed a finite element simulation of fiber network mechanical behavior to predict microscale stiffness from our macroscale measurements (**Fig. 1A**). We found that aged muscle stiffness was increased compared to young counterparts, as evidenced by a significantly reduced Green’s strain deformation (**Fig. 1B**). Using previously published formulae (19, 20), fiber mesh model predictions from experimentally-derived biaxial data revealed that aged muscle is approximately 3.5-fold stiffer than young muscle. (**Fig. 1C**).

**Figure 1.**
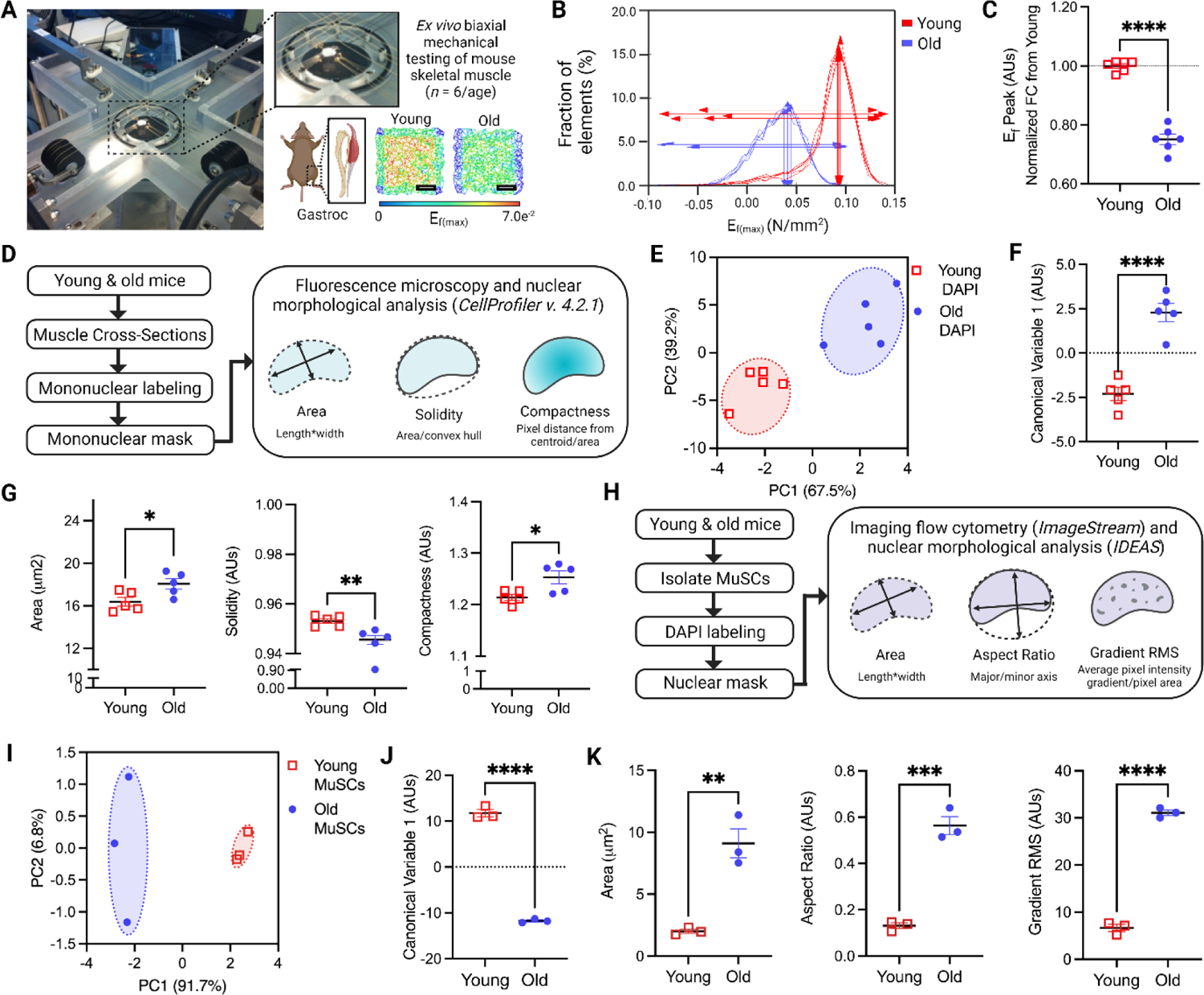
Age-related matrix stiffness is associated with aberrant MuSC nuclear morphologies. (**A**) Photograph of biaxial mechanical testing equipment. Representative heatmaps of Green’s strain energy map between young (4-6 months) and old (22–24) male mouse gastrocnemius muscle. Scale: 50 µm. (**B**) Representative histogram depicting strain deformation as a function of the fraction of elements obtained from mechanical testing (n=5/age). (**C**) Quantification of E_f(max)_ between young and old mouse muscle. (n=6/age). (**D**) Experimental workflow of nuclear morphological analysis using CellProfiler. (**E**) PCA-LDA analysis and quantification using nuclear morphological features between young and old muscle mononucleated cells (n=6/age). (**F**) Quantification of LDA canonical variable 1. (**G**) Quantification of nuclear area (μm^2^), solidity (i.e., roundness), and compactness (i.e., nuclear integrity) of mononucleated cells between young and old mouse muscle (n=5/age). (**H**) Experimental workflow of imaging flow cytometry and nuclear morphological analysis. (**I**) PCA-LDA analysis and quantification using nuclear morphological features between young and old MuSCs (n=3 mice/age group; 2,000 MuSCs represented per mouse). (**J**) Quantification of LDA canonical variable 1. (**K**) Quantification of nuclear area (μm^2^), aspect ratio (i.e., roundness), and gradient RMS (i.e., texture/granularity) of mononucleated cells between young and old muscle (n=3 mice/age group; 2,000 MuSCs represented per mouse). Unpaired student T-tests were used for Fig. C, F, G, J, K. Data are represented as mean ± SEM. *p<0.05; **p<0.01; ***p<0.001; ****p<0.0001.

Cells respond to mechanical cues through cytoskeletal remodeling, nuclear deformation, and chromosomal reorganization to modulate gene expression dictating cell fate (i.e., activation, proliferation, and/or differentiation) (21, 22). We therefore evaluated how age-related stiffness impacts the nuclear morphology of muscle mononuclear cells. To do this, we utilized *in situ* and *ex vivo* approaches to characterize nuclear morphologies using both standard fluorescence microscopy and image-based flow cytometry (ImageStream; ISX). A mask was placed over DAPI-stained mononucleated cells from muscle cross-sections, and images were assessed for a variety of nuclear morphological variables using CellProfiler™ (23) (**Fig. 1D**). Unsupervised principal component analysis (PCA) followed by supervised linear discriminant analysis (LDA) of all nuclear morphological variables (*N*=56) revealed that young and old cells display distinct morphologies, as evidenced by age-specific clustering of cells (**Fig. 1E, 1F**). Features contributing to clustering were largely attributed to significant differences in the nuclear area, solidity (i.e., roundness), and compactness (i.e., nuclear integrity) of old nuclei compared to young (**Fig. 1G**).

To corroborate the above findings with nuclear morphologies of isolated MuSCs specifically, we performed fluorescence-activated cell sorting of integrin subunit alpha 7-positive cells (*Itga7*+) followed by DAPI staining and imaging using ISX (**Fig. 1H**). PCA-LDA analyses using nuclear morphological features obtained from young and old MuSCs revealed a similar segregation of clusters according to age as compared to our *in-situ* findings (**Fig. 1I, 1J**). The primary features driving segregation of the clusters included nuclear area, aspect ratio (i.e., roundness), and gradient RMS (i.e., texture/granularity) (**Fig. 1K**). Taken together, these data demonstrate that elevated stiffness of aged muscle is associated with an altered nuclear morphology of MuSCs, *in situ* and *ex vivo*.

### Reducing matrix stiffness restores a more youthful cell state to old MuSCs

Given that nuclear architecture is a significant predictor of MuSC activation and proliferative potential (24), we hypothesized that the increased stiffness and altered nuclear morphology of old MuSCs contributes to aberrant lineage specification following activation. To test this hypothesis, we isolated and cultured young and old MuSCs on fabricated polydimethylsiloxane (PDMS) substrates calibrated to approximate the physiological stiffness of young (soft; 12 kPa) and old (stiff; 29 kPa) muscle, after which we performed morphometric analysis using ISX (**Fig. 2A**). A combination of 8 nuclear morphological variables were used as input for PCA-LDA analyses, which demonstrated no stiffness-related clustering of young MuSCs cultured on either soft or stiff substrates (**Fig. 2B, 2C**). In contrast, old MuSCs cultured on a soft, but not a stiff, substrate showed tight clustering (**Fig. 2D, 2E**). Interrogation of variables contributing to LDA analyses revealed there were no differences in young MuSCs for any of the measured variables, regardless of matrix condition. In contrast, LDA was significantly different when comparing old MuSCs seeded onto soft versus substrates (**Fig. 2E**). Accordingly, we observed a significant reduction in the nuclear area in old MuSCs on soft substrates compared to stiff (**Fig. 2F, 2G**).

**Figure 2.**
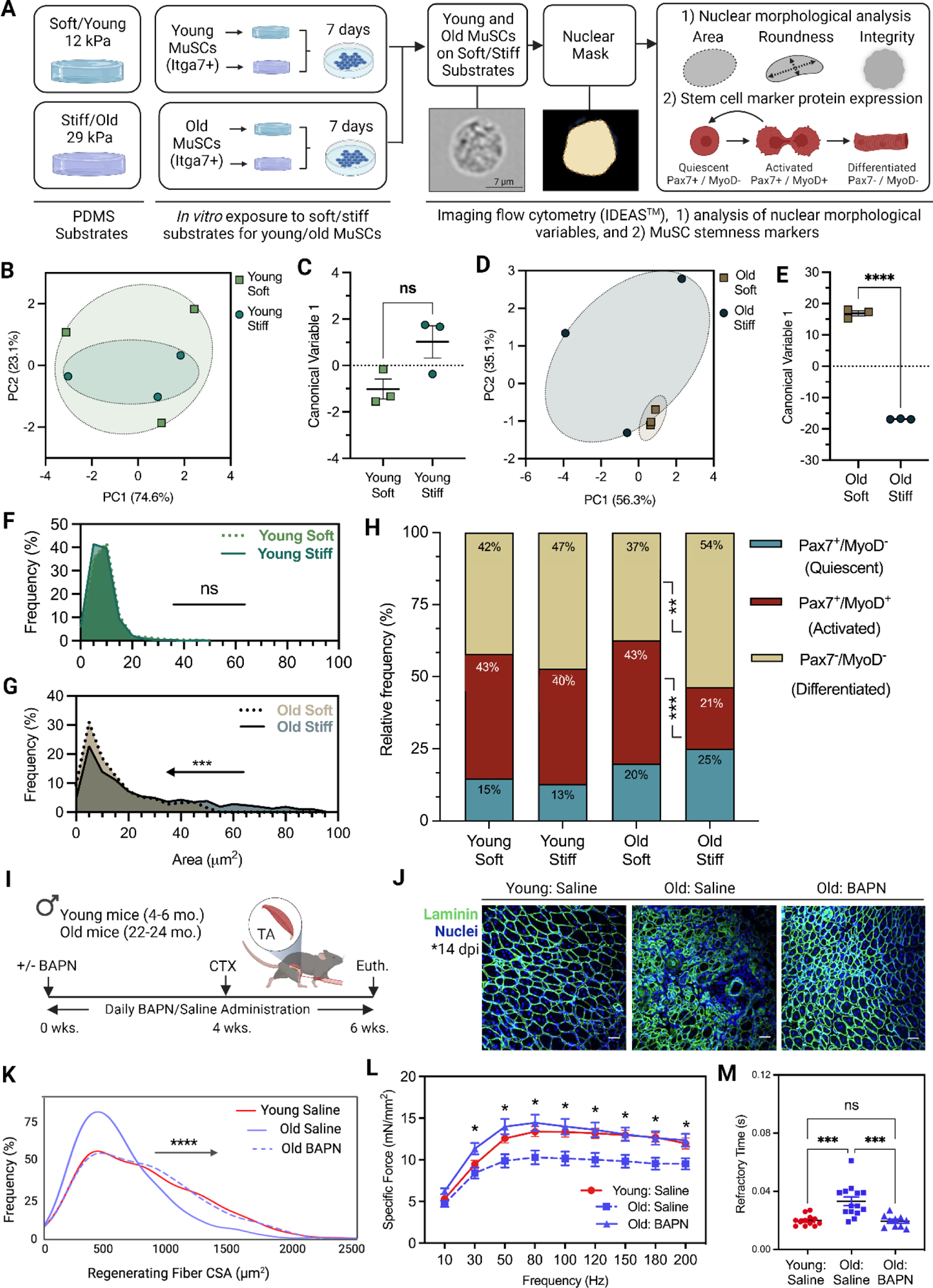
Reducing matrix stiffness restores a more youthful cell state to old MuSCs. (**A**) Experimental design of PDMS fabrication, MuSC culture, and ISX analyses. (**B, C**) PCA-LDA analysis and quantification using nuclear morphological features between young MuSCs cultured on either soft or stiff substrates (n=3 mice/age group; 2,000 MuSCs represented per mouse). (**D, E**) PCA-LDA analysis and quantification using nuclear morphological features between young MuSCs cultured on either soft or stiff substrates (n=3 mice/age group; 2,000 MuSCs represented per mouse). (**F**) Relative frequency of young MuSC nuclear area (μm^2^) on either soft or stiff substrates. (**G**) Relative frequency of old MuSC nuclear area (μm^2^) on either soft or stiff substrates. (**H**) Relative frequency of MuSCs with positive fluorescent signals for Pax7 and MyoD between young and old MuSCs cultured on either soft or stiff substrates. Pax7+/MyoD-: quiescent, Pax7+/MyoD+: Activated, Cycling, Pax7-/MyoD-: Differentiated. (**I**) Experimental design of β-aminopropionitrile (BAPN) injections for preconditioning of skeletal muscle prior to injury between BAPN and saline control injected mice (n=5-6 mice/group). (**J**) Representative images of muscle cross sections immunolabeled with laminin (green) and DAPI (nuclei) between young mice injected with control saline solution and old mice with either saline or BAPN injections. Scale bar: 200 μm^2^ (**K**) Frequency distribution of regenerating muscle fiber area (μm^2^) of TA muscle 14-days post-injury (dpi) from BAPN and saline-treated old mice. (n=12-14 mice/group) (**L**) Force-frequency curves from *in situ* contractile testing of recovering TA muscles 14 dpi between BAPN and saline-treated old mice. (**M**) Half relaxation time from in situ contractile testing of recovering TA muscles 14 dpi between BAPN and saline-treated old mice. Unpaired student T-tests were used for Fig. C, E. Nonparametric Kolmogorov-Smirnov tests were used for Fig. F, G, K. ANOVAs were used for Fig. H, L, M, followed by Tukey’s post hoc when appropriate. Data are represented as mean ± SEM. *p<0.05; **p<0.01; ***p<0.001; ****p<0.0001.

To determine whether changes in the nuclear morphology of old MuSCs on soft substrates are accompanied by changes in cell state markers, we next performed ISX of MuSC lineage markers corresponding to quiescence (Pax7+/Myod-), activation and/or cycling (Pax7+/MyoD+), and differentiation (Pax7-/MyoD-). As was the case for morphological metrics, we observed no differences in relative marker expression in young MuSCs on soft versus stiff matrices (**Fig. 2H**). However, old MuSCs were highly sensitive to matrix condition, as soft substrates increased the percentage of activated/cycling old MuSCs compared to cells cultured on stiff substrates (43% vs. 21%, respectively; **Fig. 2H**). The fraction of differentiated MuSCs (Pax7-/MyoD-) was also reduced when old MuSCs were cultured on soft substrates as compared to a stiff substrate (37% vs. 54%, respectively; **Fig. 2H**). These data suggest stability in the young MuSC phenotype, regardless of substrate stiffness. Old MuSCs, however, are highly responsive to extrinsic biophysical cues, whereby reduced matrix stiffness promoted an increased number of activated and cycling cells.

To explore the potential physiological relevance of the above findings, we next pharmacologically manipulated the mechanical properties of skeletal muscle in aged mice using β-aminopropionitrile (BAPN). BAPN is a chemical inhibitor of lysyl oxidase (LOX), an enzyme responsible for the age-related accumulation of crosslinked collagen and elevated tissue stiffness, not only in skeletal muscle (9), but also in the lungs, heart, and bone (25–28). We performed daily subcutaneous injections of BAPN for a total of six weeks in young and old mice. Four weeks following the start of BAPN injections, animals received cardiotoxin injections bilaterally in tibialis anterior muscles (**Fig. 2I**). We hypothesized that decreased matrix cross-linking and, by extension, matrix stiffness (29), would improve the regenerative capacity of aged muscle following an injury. Two weeks after recovery from injury, aged mice that received BAPN exhibited improved regeneration, evidenced by increased regenerating fiber cross-sectional area (CSA), increased force producing capacity, and a reduced refractory period following stimulation (**Fig. 2J-2M**). Collectively, these results corroborate our *in vitro* findings and demonstrate that manipulation of the aged MuSC microenvironment benefits functional skeletal muscle regeneration *in vivo*.

### Post-transcriptional dynamics contribute to stiffness-mediated lineage specification

Thus far, our results have shown that reducing substrate stiffness impacts the activated and cycling fraction of old MuSCs, which may be reflective of enhanced self-renewal. As such, we sought to better understand the molecular mechanisms constituting these changes. To do this, we performed single cell RNA sequencing (scRNA-seq) and *in silico* lineage analysis of young and old MuSCs on soft and/or stiff substrates. Following cell/transcript quality control and filtration of technical artifact (**Fig. S1A-E**), dimensionality reduction and trajectory analyses of MuSCs were performed using Monocle 3.0 (30) (**Fig. 3A**). Monocle 3.0 is a lineage analysis algorithm that organizes cells based on cell state to identify cellular lineage progression (30, 31). Dimensionality reduction using PCA and UMAP visualization of MuSCs showed no differences in clustering as compared to Monocle 3.0 trajectory projections (**Fig. SF-H**). Therefore, we focused our subsequent analyses of MuSCs using Monocle 3.0 alone.

**Figure 3.**
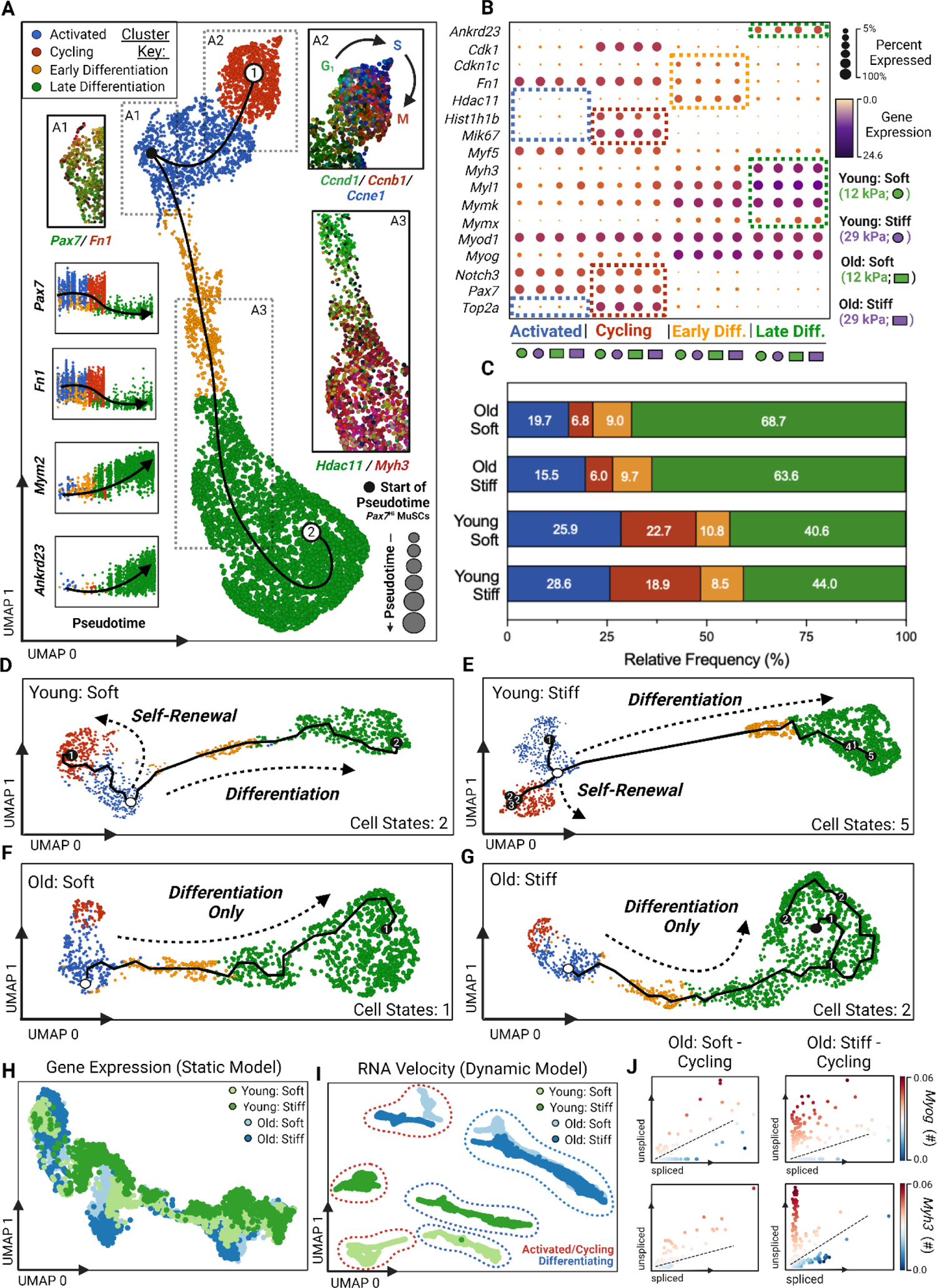
Post-transcriptional dynamics contribute to stiffness-mediated lineage specification. (**A**) UMAP representation of aggregated MuSCs used for trajectory analysis from young/old and soft/stiff conditions. Primary plot is colored based on stage of lineage progression (Activated: blue, Cycling: red, Early Differentiation: green, Late Differentiation: orange). Pseudotime and lineage progression is indicated by bubble size, increasing from Activated to Cycling and Differentiated cell states. Expression of fibronectin (**A1**; *Pax7*/*Fn1*; activated), cyclin D1 (**A2**; *Ccnd1*; cycling), cyclin B1 (**A2**; *Ccbn1*; cycling), cyclin E1 (**A2**; *Ccne1*; cycling), histone deacetylase 11 (**A3**; *Hdac11*; early differentiation), and embryonic myosin heavy chain (**A3**; *Myh3*; late differentiation) is provided via dotted insets for validation of cluster classification. Correlation plots showing expression of *Pax7* (Activated, Cycling), *Fn1* (Activated, Cycling), *Mym2* (Early, Late Differentiation), and *Ankrd23* (Early, Late Differentiation) changing as a function of pseudotime is also provided. (**B**) Bubble plot indicating cluster-specific gene expression. Bubble size corresponds to the percentage of cells within each subpopulation expressing the gene of interest, and average expression value is indicated by heatmap. Dotted boxes indicate subpopulation-specific gene expression (i.e., *Cdk1* is expressed only in the Cycling subpopulation of MuSCs) (**C**) Relative frequency of MuSCs per lineage between young and old MuSCs on soft and stiff matrices. (**D-G**) Lineage specification analysis using Monocle 3.0 for young and old MuSCs exposed to soft and/or stiff substrates, each colored by subpopulation (Activated: blue, Cycling: red, Early Differentiation: green, Late Differentiation: orange). (**H**) Dimensionality reduction method, UMAP, for visualization of the four samples using gene expression. (**I**) UMAP for visualization of the four samples using RNA velocity. (**J**) Phase portraits of spliced-to-unspliced gene counts for myogenin (*Myog*) and embryonic myosin heavy chain (*Myh3*) between old cycling and activated MuSCs cultured on soft and stiff substrates.

Analyzing MuSCs from all four groups in aggregate (i.e., young/old, and soft/stiff MuSCs), trajectory analyses identified a myogenic continuum originating from MuSCs with the highest expression of MuSC marker, *Pax7*, and then bifurcating towards either a self-renewing or a differentiating trajectory (**Fig. 3A, S2I**). Following unbiased graph-based clustering, a total of four distinct subpopulations of MuSCs were identified along each trajectory, which we labeled “Cycling” and “Activated” as part of the self-renewing trajectory, and “Early Differentiation”, and “Late Differentiation” as part of the differentiating trajectory (**Fig. 3A, 3B**). The validity of subpopulation classification was demonstrated by high expression of genes that have been previously shown to be indicative of each cell state (i.e., Activated-*Fn1*^Hi^, Cycling-*Ccnd1*^Hi^/Ccnb1^Hi^/Ccne1^Hi^, Early Differentiation-*Myog*^Hi^/Hdac11^Hi^, and Late Differentiation-*Myh3*^Hi^; (32–35)) (**Fig. 3A1-3A3, 3B**). Moreover, self-renewing genes were highly expressed towards the start of pseudotime, while differentiating genes were highly expressed by the end of pseudotime. When cells were segregated according to group (i.e., young versus old on soft versus stiff), trajectory analysis revealed that, as expected, young MuSCs displayed both self-renewing and differentiating trajectories, regardless of the substrate stiffness (**Fig. 3C-3E**), though a stiff substrate did introduce a bifurcation in each direction. That is, we observed an additional branch in both the self-renewing and differentiation trajectories (**Fig. 3E**). Such bifurcations are indicative of increased complexity of the parent trajectory (30). In contrast, we found that old MuSCs displayed relative transcriptional rigidity, adopting only a single lineage towards differentiation on both soft and stiff substrate conditions (**Fig. 3C, 3F, 3G**). The number of cycling old MuSCs, while not altogether absent, was insufficient to constitute a bona fide self-renewing trajectory, regardless of whether cells were seeded onto a soft or stiff substrate (**Fig. 3C, 3F, 3G**). Similar to lineage analyses using Monocle 3.0, there were no significant differences in differential gene expression for old MuSCs seeded on soft versus stiff substrates (**Fig. S2K)**. We found this observation surprising given that old MuSCs seeded onto soft substrates displayed a phenotype indicative of enhanced self-renewal when compared to old cells seeded on a stiff substrate (**Fig. 2H**).

The above findings suggest a disconnect between morphological/protein level changes and those at the transcriptional level. This is consistent with previous observations showing that steady-state *(static)* measures of RNA abundance often fail to correlate with protein levels (36–38). As an alternative, a growing number of studies have demonstrated that post-transcriptional gene regulation plays an underappreciated role in shaping the proteome (39). We therefore posited that non-steady-state *(dynamic)* RNA metrics may better capture the transcriptional changes mediating the phenotypic responses of old MuSCs on a soft matrix. RNA velocity provides such a metric by accounting for the relationship between the relative abundance of both unspliced and spliced RNA, thereby providing estimated rates of splicing and degradation (40). As evidence of the distinct features captured by dynamic versus static approaches, PCA revealed only a single cluster when considering static gene counts, indicating few differences across the four experimental groups (**Fig. 3H**). In contrast, RNA velocity yielded a total of six clusters corresponding to cell state, age, and matrix condition (**Fig. 3I**). Post-transcriptional regulators DEAD-box helicase 27 (*Ddx27*) and ubiquitin specific peptidase 39 (*Usp39*) serve as just two examples of such a distinction between static versus dynamic assessments (**Fig. S2L)**.

When considering RNA velocity, old MuSCs on soft versus stiff substrates diverged in the activated and cycling cluster of cells, while the differentiating group of cells on each substrate condition were overlapping (**Fig. 3I**). To investigate whether cluster segregation was the result of impaired old MuSC self-renewing potential and a predisposition towards differentiation, we inferred the future states of cycling and activated old MuSCs using phase portraits. Phase portraits delineate whether transcripts are considered to be in steady-state, increased velocity (i.e., an increased abundance of unspliced to spliced transcripts), or decreased velocity (i.e., an increased abundance of spliced to unspliced transcripts) (40). Phase portraits for early differentiation marker, myogenin (*Myog*), and late differentiation marker, *Myh3*, revealed increased velocity for each marker in old MuSCs on stiff substrates (**Fig. 3J**). Old MuSCs on soft substrates, however, showed relatively few unspliced or spliced transcripts of either marker, suggesting that soft matrices maintain a self-renewing state for old MuSCs. One interpretation of these results is that, while static modeling identified negligible differences in old MuSC trajectories when considering soft versus stiff substrates, dynamical modeling revealed that stiff substrates predispose old MuSCs towards differentiation while soft substrates preserve a self-renewing state. These data raised the hypothesis that the effect of substrate stiffness on old MuSC lineage progression may be primarily mediated through post-transcriptional mechanisms.

### Soft matrices reduce RNA decay pathway activity to preserve self-renewal in old MuSCs

To further investigate lineage specification trajectories using RNA velocity, we leveraged *dynamo*, a robust machine learning framework that utilizes RNA velocity of the cell’s transcriptome to reconstruct continuous velocity vector fields (41). *Dynamo* not only provides insight into the future state of cells through visualization of RNA velocity vector fields, but it also uses differential geometry analysis to identify the “attractors” that govern cell gravitation towards state-specific stable points. Attractors represent nodes to which cells converge along the trajectory, thereby reflecting deterministic lineage specification tendencies. Using this approach, we evaluated whether reducing matrix stiffness influences the stability and/or presence of “attractors” related to self-renewal in old MuSCs (**Fig. 4A**).

**Figure 4.**
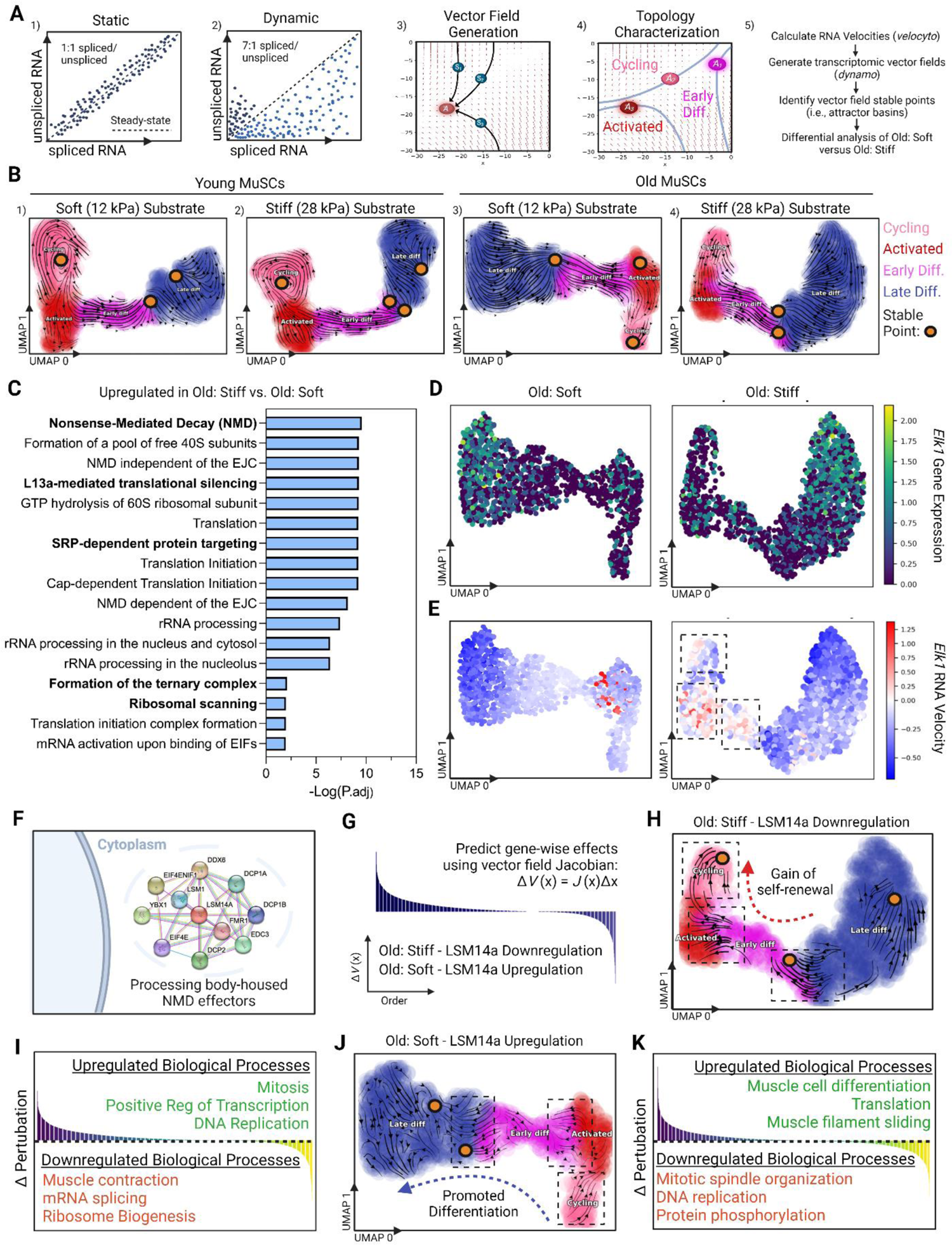
Soft matrices reduce RNA decay pathway activity to preserve self-renewal in old MuSCs. (**A**) Schematic of transcriptional vector field generation and workflow. (**A1**) Example of a gene of interest (GOI) assumed to be in steady state when only mature gene counts are used. (**A2**) Example of a GOI in non-steady-state conditions, identified using the ratio of spliced to unspliced transcripts (i.e., RNA velocity). (**A3**) Example of a transcriptomic vector field generated using the RNA velocity values of each gene. (**A4**) Topology characterization of the vector field. Separatrices (blue lines) identify the boundaries of attractor basins (*A*_1-3_) corresponding to each stable cellular state. (**A5**) Summary of experimental workflow using vector fields to identify how soft and stiff matrices influence self-renewal in old MuSCs. (**B**) Velocity vector fields depicting cellular fate progressions on UMAP space for young (**B1-2**) and old (**B3-4**) MuSCs on soft or stiff matrices. Coloring denotes stage of lineage progression. Stable fixed points indicate local stable phenotypes to which MuSCs are attracted. (**C)** Functional gene set enrichment of unique genes in rank-order identified using random forest classification between old MuSCs on soft and stiff matrices. Processes associated with the cellular stress response and RNA decay are highlighted via bolded text. (**D**) Comparison of *Elk-1* gene expression between old MuSCs on soft and stiff substrates. (**E**) Comparison of *Elk-1* RNA velocity between old MuSCs on soft and stiff substrates. Dotted boxes indicated elevated *Elk-1* RNA velocity in the cycling and activated states for old MuSCs on stiff matrices. (**F**) Schematic of physical interaction networks between processing body marker LSM14a mRNA processing body assembly factor (*LSM14a*) and NMD protein machinery. LSM14a physical interactions shown using STRING with interaction confidence set to high (0.7 out of 1.0). (**G**) Vector field perturbation scheme. (**H)** *In silico* perturbation of old MuSCs on stiff matrices of downregulated *LSM14a*. Cluster colors correspond to stage of lineage progression. Only significantly affected vectors are shown. (**I**) Gene set enrichment analyses of biological processes up and downregulated as a result of *LSM14a* downregulation in old MuSCs exposed to stiff matrices. (**J**) *In silico* perturbation of old MuSCs on soft matrices of upregulated *LSM14a*. Cluster colors correspond to stage of lineage progression. Only significant vectors are shown. (**K**) Gene set enrichment analyses of biological processes up and downregulated as a result of LSM14a upregulation in old MuSCs exposed to soft matrices. A Benjamini-Hochberg false discovery rate of 0.05 was used for multiple testing correction, and gene term size limits between 5-500 were used for gene set enrichment analyses (Fig. C, I, and K).

Following vector-field generation, we observed numerous single cell trajectories (indicated by vector orientation) for young and old MuSCs on each substrate condition, with attractors identified in various intermediate cell states (**Fig. 4B1-4**). In contrast to static expression-based approaches, *dynamo* revealed age-, substrate-, and subpopulation-dependent stable points (attractors) for both young and old MuSCs on each substrate condition (**Fig. 4B1-4**). Young MuSCs exposed to soft versus stiff substrates demonstrated similar attractor profiles, with subpopulations that spanned both self-renewing and differentiation trajectories (**Fig. 4B1-2**). These analyses are consistent with our previous findings demonstrating the resilience of young MuSCs even when exposed to a stiff matrix (**Fig. 3E**). The attractors for old MuSCs on a stiff substrate indicated a predisposition of towards differentiation (**Fig. 4B4**). However, on a soft substrate, new stable points in the activated and cycling states of old MuSCs emerged, suggesting enhanced self-renewal (**Fig. 4B3**).

To identify genes driving the attraction of old MuSCs seeded on a soft, but not stiff, matrix towards self-renewing intermediate cell states, we performed random forest classifications (42). Random forests identified a discrete gene set with corresponding differential RNA velocities between the two groups. Functional gene set enrichment analyses summarized these genes to be highly associated with nonsense mediated RNA decay (NMD) and the cellular stress response (43–46) (**Fig. 4C**). The velocities of nearly all identified genes were reduced in old MuSCs on soft matrices relative to stiff matrices, suggesting that an aged microenvironment enhances effectors of stress and RNA decay (**Fig. S2A**). To determine potential transcription factors mediating the activation of the stress response and RNA decay in old MuSCs on stiff matrices, we overlaid the gene set identified above with a curated database of known transcription factor (TF) – target gene pairs using TRANSFAC (47). The TF most associated with the identified genes was ETS Like-1 protein Elk-1 (*Elk1*), a member of the stress-activated protein kinase superfamily and relative of p38 mitogen-activated protein kinase (MAPK) (**Fig. S2B**) (45). P38 MAPK is a known contributor to the loss of self-renewal and stemness in old MuSCs (48). It is worthwhile to note that there were no differences in *Elk1* when considering static gene expression levels between old MuSCs exposed to either soft or stiff matrices (**Fig. 4D**). In contrast, *Elk1* displayed increased velocity at the boundary between activated and differentiating lineages for old MuSCs (**Fig. 4E**). Given increased velocity suggests an increased rate of splicing and, therefore, a greater number of mature transcripts, these findings suggest a potential relationship between matrix stiffness, the cellular stress response, and subsequent changes in cell fate. Importantly, there was an overall decrease in *Elk1* RNA velocity in old MuSCs on soft substrates compared to stiff, implying reduced activation of cellular stress and RNA decay (**Fig. 4E**). Together, these data suggest that the emergence of stable points in states related to self-renewal for old MuSCs on a soft substrate may be attributed to decreased stress activation and RNA degradation.

Does RNA decay mediate the effects of age-related matrix stiffness on old MuSC fate? To answer this question, we applied the perturbation feature in *dynamo,* which leverages the vector field Jacobian. This approach allows genes of interest (GOIs) to be computationally tuned to observe how the velocities of the remaining genes change, and subsequently how lineage progression is affected. In this way, we can determine how GOIs control lineage specification trajectories of the MuSC population. This *in silico* approach was therefore used to test our hypothesis that stiff matrices increase RNA degradation and promote differentiation of old MuSCs, while soft matrices reduce RNA decay to preserve self-renewal (**Fig. 4G**). Since NMD machinery is a complex arrangement of proteins that have redundant roles in RNA degradation, we manipulated expression of a foundational NMD scaffold protein, LSM14a mRNA processing body assembly factor (*LSM14a*), which is responsible for housing and protecting NMD effectors (49) (50, 51) (**Fig. 4F**). We hypothesized that computational inhibition of *LSM14a* in old MuSCs on stiff matrices would decrease RNA decay and restore self-renewal, thus inducing a more youthful lineage trajectory profile. Conversely, we hypothesized that computational overexpression of *LSM14a* in old MuSCs on soft matrices would reinforce vectors towards differentiation, thereby abrogating the beneficial effect of a soft matrix on old MuSC self-renewal (**Fig. 4G**).

*In silico* downregulation of *LSM14a* caused the trajectories of old MuSCs cultured on stiff matrices to reverse vector fields, predisposing cells away from differentiation in favor of an undifferentiated, self-renewing state (**Fig. 4H**). Characterization of genes upregulated by computational inhibition of *LSM14a* revealed an enrichment of biological processes related to cellular division and mitosis, including increased expression of self-renewal genes such as H2a histone family member Z (*H2afz*) and syndecan-4 (*Sdc4*) (52–54) (**Fig. 4I**). Biological processes downregulated as a result of reduced *LSM14a* expression were related to muscle contraction, with corresponding reduced expression of myosin heavy chain gene isoforms including −1, −3, and −7 (*Myh1, Myh3, Myh7*, respectively) (**Fig. 4I**). On the other hand, *in silico* upregulation of *LSM14a* in old MuSCs on soft matrices altered the direction of vectors away from a cycling, activated, and early differentiated state and towards a fully differentiated one (**Fig. 4J**). Consistent with this overall trend, characterization of biological processes up- and down-regulated with *LSM14a* upregulation revealed enrichment of processes associated with muscle cell differentiation and mitosis, respectively (**Fig. 4K**). To rule out the possibility that these findings are age-dependent, we performed the same perturbations in young MuSCs. We observed that upregulation of *LSM14a* in young MuSCs on soft matrices had similar effects as were observed in old cells, that is an aversion from self-renewing trajectories towards differentiation (**Fig. S2C**). Taken together with protein and morphological-level assessments, these results demonstrate that age-related increases in matrix stiffness stimulate activation of RNA decay pathways, predisposing MuSCs towards differentiation at the expense of self-renewal.

## Discussion

Here, we provided evidence that age-related increases in matrix stiffness compromise MuSC self-renewal through stimulation of RNA decay, and that these effects can be reverted by exposing old MuSCs to a soft matrix. We showed that aged muscle is approximately 3.5-fold stiffer than young muscle, alterations we found to be associated with aberrant nuclear morphologies in old MuSCs. Using substrates engineered to mimic the biophysical properties of young versus aged muscle, we next found that aging affected the sensitivity of MuSCs to extrinsic biophysical signals. That is, whereas an increase in matrix stiffness had little effects on the phenotype of young MuSCs, old MuSCs exposed to a young-like matrix displayed a more youthful phenotype, as evidenced by an increased number of old cells in the cycling and activated state (i.e., self-renewing cells). When we compared static versus dynamic analyses of single cell lineage trajectories, we found that dynamic assessments using RNA velocity captured the beneficial effect of soft matrices on old MuSC responses. Specifically, a soft matrix reduced RNA degradation and the stress response in old MuSCs, resulting in the promotion of a self-renewing subpopulation. Our results highlight previously unrecognized mechanisms of old MuSC self-renewal in response to extrinsic biophysical cues, while also suggesting rejuvenation of the cellular milieu as a strategy to restore youthful fate transitions in old MuSCs.

The role of matrix stiffness on old MuSC fate has been investigated in the context of both differentiating and self-renewing MuSCs. We and others have demonstrated the numerous alternative fates old MuSCs adopt when exposed to age-related matrix stiffness, evidenced by fibrogenic, inflammatory, and senescent conversions, ultimately contributing to hampered myofiber repair in aged skeletal muscle (2, 8, 55, 56). Reports investigating the effects of stiffness on old MuSC self-renewal, however, are fewer in number. Cosgrove et al. demonstrated enhanced self-renewal of old MuSCs through exposure to p38 MAPK pathway inhibition combined with culture on soft hydrogels (57). Similarly, our computational analyses identified that aged matrices stimulated stress activated MAPK signaling, which corresponded with a loss of self-renewing potential in old MuSCs. While aging was not studied directly, Moyle et al. recently showed that matrix stiffness synergizes ligands essential for self-renewal, and further went on to postulate that age-related stiffness may directly contribute to lost regenerative capacities with age (58). However, mechanisms as to how aged matrices and kinase signaling disrupt the ability of old MuSCs to self-renew have remained unclear. Our current work demonstrates that aged matrices stimulate cell stress pathway activation and subsequently RNA decay to degrade the transcripts essential for self-renewal, driving old MuSCs towards differentiation.

The relationship between RNA degradation and cellular fate transitions has been documented in rodents (59–63), fruit flies (64), flatworms (65), and humans (66–68), spanning embryonic (66–68) and adult stem cells (63, 68). The post-transcriptional control of MuSC fate has also been reported (59, 60, 63, 69). Matrix stiffness is a well-documented morphogen that dictates fate transitions in many cell types (5, 8, 70–72), including MuSCs (5, 8), and the results from our current studies implicate a previously unestablished link between matrix stiffness and RNA decay in MuSCs that contributes to age-related alterations in lineage specification.

Increased matrix stiffening, a quintessential feature of aged tissues and regarded by some as a hallmark of aging (73), can in many ways be considered a form of cellular stress. Like many forms of stimuli in cell signaling, a “sweet spot” exists, in that too much or too little stress causes, at best, enhanced cellular resilience (74), and at worst, pathogenic outcomes such as cell death (75). As matrix stiffness is a primary driver of mechanotransductive signaling, when increased matrix stiffness of aged tissues falls outside a range of beneficial signaling, it is instead received by the cell as a form of pathogenic stress. Cells respond to stress through evolutionarily conserved surveillance mechanisms, which reorient both transcriptional and translational processes with the goal of re-establishing homeostasis (76). Remodeling is achieved, at least in part, by stalling protein synthesis and degrading transcripts related to the source of stress (77, 78). For example, during endoplasmic reticulum (ER) stress and the unfolded protein response (UPR), not only is protein synthesis halted as a result of the stress response, but the NMD pathway is also activated to degrade already transcribed mRNAs in order to rapidly terminate the UPR (79). In terms of matrix stiffness, it has recently been shown that pathological stiffness initiates DNA damage and activates the DNA damage repair pathway (DDR) (80). Upon activation of the DDR, both ribosome stalling and activation of the NMD pathway are initiated (81, 82) to clear repair response-related mRNAs and terminate the DDR. In our studies, we identified Elk1-mediated ternary complex formation, translational silencing, and NMD activation as primary drivers of the differential responses of old MuSCs when seeded onto stiff versus soft matrices. Considering Elk1 is a stress-activated MAPK (45) coupled with numerous studies demonstrating aberrant MAPK signaling in old MuSCs (48, 83, 84), our results suggest that the inability to respond to stress in old MuSCs contributes to an impaired self-renewing state. Future studies directly investigating the role of NMD during stress and stem cell fate transitions, particularly through the lens of aging stem cells, may yield intriguing insights into the loss of potency in old MuSCs.

To better understand the role of NMD in the sensitivity of old MuSCs to substrates of varying stiffness, we performed *in silico* manipulation of *LSM14a*. LSM14a is a structural protein necessary for processing body (P-body) formation, a cytoplasmic storage site for mRNA degradation and, reported recently, a gatekeeper for cell fate transitions (68, 85, 86). In human embryonic stem cells as well as adult progenitors (including myoblasts), P-body ablation resulted in release of mRNAs encoding for pluripotent-specific transcription factors which, when translated, induced a hyper-pluripotent state (68). While the impact of aging on P-bodies was not investigated in this study, the predominance of biological processes associated with NMD in stem cells, together with the role of LSM14a observed in our studies, suggests that changes in P-body dynamics with age may impair the ability of MuSCs to self-renew upon division. Moreover, these data further suggest that age-related matrix stiffness may impact P-body dynamics, potentially influencing the lineage specification of old MuSCs. The potential interaction of P-body dynamics and mechanical signals from the stem cell microenvironment is currently an unexplored area of research. Further research on these topics, as well as whether and how these mechanisms can be extrapolated to other adult progenitors in aged individuals, is needed.

The finding that young MuSCs maintain pluripotency regardless of substrate condition, while old MuSC lineage trajectories are readily manipulated by soft substrates is consistent with observations of a youthful resilience in the face of perturbation that is lost with increasing age (1). Over thirty years ago, principles of nonlinear dynamics and chaos theory were first used to describe the loss of complexity with aging that contributes to an inability of aged systems to respond to stress (87). It was postulated that young systems, spanning molecular to organismal scales, maintain vitality through a combination of highly complex and finely tuned biological processes that are required to efficiently respond to stress and maintain homeostasis. With aging, the loss of complexity underlies the inability of a system to respond to physiologic stress, causing functional breakdown. An example of these concepts at the organ level is seen in heart rate (HR) variability with aging. HR variability is highly irregular in young individuals, but variability drops considerably in old individuals (87). This loss of variability (i.e., complexity) renders older adults less able to adapt to stress, which associates with an increased likelihood of cardiac pathology. Similarly, in our study, young MuSCs were robust to perturbation even in the face of an aged matrix, showing minimal change in transcriptomic and phenotypic metrics. In contrast, the low transcriptional complexity of old MuSCs (i.e., impaired regulatory mechanisms) is evidenced by an inability to maintain self-renewal when challenged with the stress of a stiff matrix. The consequence of low complexity is evidenced by increasing rates of clonal drift in aging stem cell compartments, including both hematopoietic stem cells (88) and MuSCs (89). Importantly, we found that the consequence of low resilience in old MuSCs was mitigated when the source of stress (i.e., aged matrix stiffness) was minimized, as evidenced by an increased self-renewing potential. Just as nonlinear system approaches were required to describe the irregularity of aged heart rate variability, only *dynamo* RNA velocity vector fields, also derived from nonlinear mathematical approaches, effectively captured the transcriptional dynamics governing cell fate (41). Collectively, our results implicate old MuSC self-renewal as a system that is subject to a loss of complexity, and that aged matrix stiffness is a source of cellular stress.

### Limitations of Study

There are limitations of the current findings worth noting. First, only two age groups were studied, young and old, thereby limiting a holistic understanding of how the principles identified here applies to the process of aging over the lifespan. Recent work from our group and others have shown that aging is a non-linear process (90, 91), and additional ages will likely provide novel insights into whether there is a tipping point at which MuSC display altered sensitivity to extrinsic matrix cues, or if the decline is gradual and progressive (90). Second, only male mice were used. Future studies are needed to compare the trajectory of stem cell fate changes over time and according to sex. Third and finally, while our understanding of post-transcriptional control over cellular function is growing rapidly, most research in this area is limited by snapshot-based measurements. Recent reports have shown large fractions of the transcriptome never undergo translation, thereby confounding our understanding of the relationship between gene expression and protein production (92, 93). For instance, single cell metabolic labeling of nascent RNA followed by fractionization of subcellular compartments revealed that only a subset of transcribed RNAs get exported from the nucleus, while a large fraction gets degraded (93). Moreover, while the importance of spatial information has gained increased appreciation since the advent of spatial scRNA-seq, recent studies have shown a spatial distribution of the transcriptome and the pronounced effect on rates of protein production, for which current sequencing technologies do not account (92, 94). These limitations represent interesting opportunities for future investigation.

## Acknowledgements

The authors thank Samuel K. Luketich and Drake D. Pedersen for their help with biaxial mechanical testing and Philip R. LeDuc for guidance with PDMS fabrication. All figures were prepared using Biorender. Data preprocessing and analysis were performed using Partek Flow Genomics Suite.

## Author Contributions

Author contributions were determined using the CRediT model: Conceptualization: ZH, SH, HM, HI, AS, KW, GG, DV, TR, FA. Methodology: ZH, SH, HM, HI, AS, GG, AM, GN, AD. Investigation: ZH, SH, HM, HI, AS, AM, GN, AD. Visualization: ZH, SH, JX, FA. Funding acquisition: JX, FA. Project administration: FA. Supervision: JX, FA. Writing – original draft: ZH, FA. Writing – review & editing: ZH, SH, HI, AS, KW, GG TR, JX, FA.

## Supplemental Figures

### Supplemental Figure Legends

**Figure S1.**
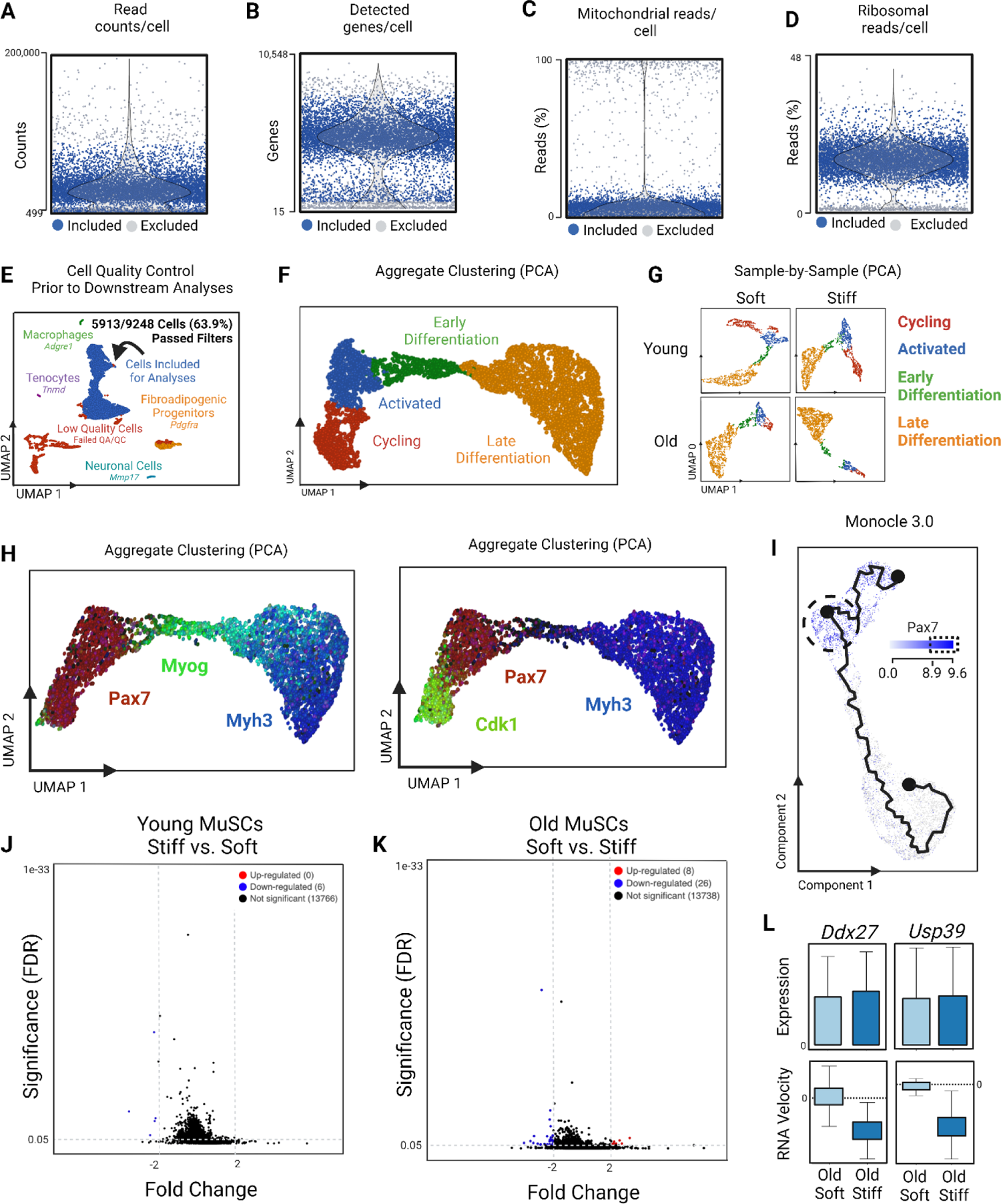
Single cell RNA sequencing data preprocessing of low-quality cells using metrics of: (**A**) read counts per cell, (**B**) detected genes per cell, (**C**) mitochondrial reads per cell, and (**D**) ribosomal read counts per cell. Grey colored cells outside the upper and lower quartiles of each metric were excluded while blue colored cells were included for analyses. (**E**) UMAP visualization of cells that passed quality control analyses but were computationally identified not to be a homogenous pool of MuSCs, likely from the small amount of error occurring from FACS sorting. Trace populations of fibro-adipose progenitor cells, macrophages, tenocytes, and neuronal cells were excluded to prevent confounders. (**F**) Dimensionality reduction using PCA projected onto UMAP space of all samples in aggregate. (**G**) Dimensionality reduction using PCA projected onto UMAP of each individual sample. (**H**) PCA-derived UMAP of demarcating genes used for cluster classification: *Pax7*, cyclin dependent kinase 1 (*Cdk1*), *Myog*, and *Myh3*. (**I**) UMAP projection of trajectory analysis colored by *Pax7*-Hi cells, representing the start of pseudotime progression. Volcano plot of differentially expressed genes between (**J**) young, and (**K**) old MuSCs on soft versus stiff matrices (−2>FC>2; p.adj<0.05). (**L**) Comparison of gene expression and RNA velocity of DEAD-box helicase 27 (*Ddx27*) and Ubiquitin Specific Peptidase 39 (*Usp39*) between old MuSCs on soft and stiff matrices.

**Figure S2.**
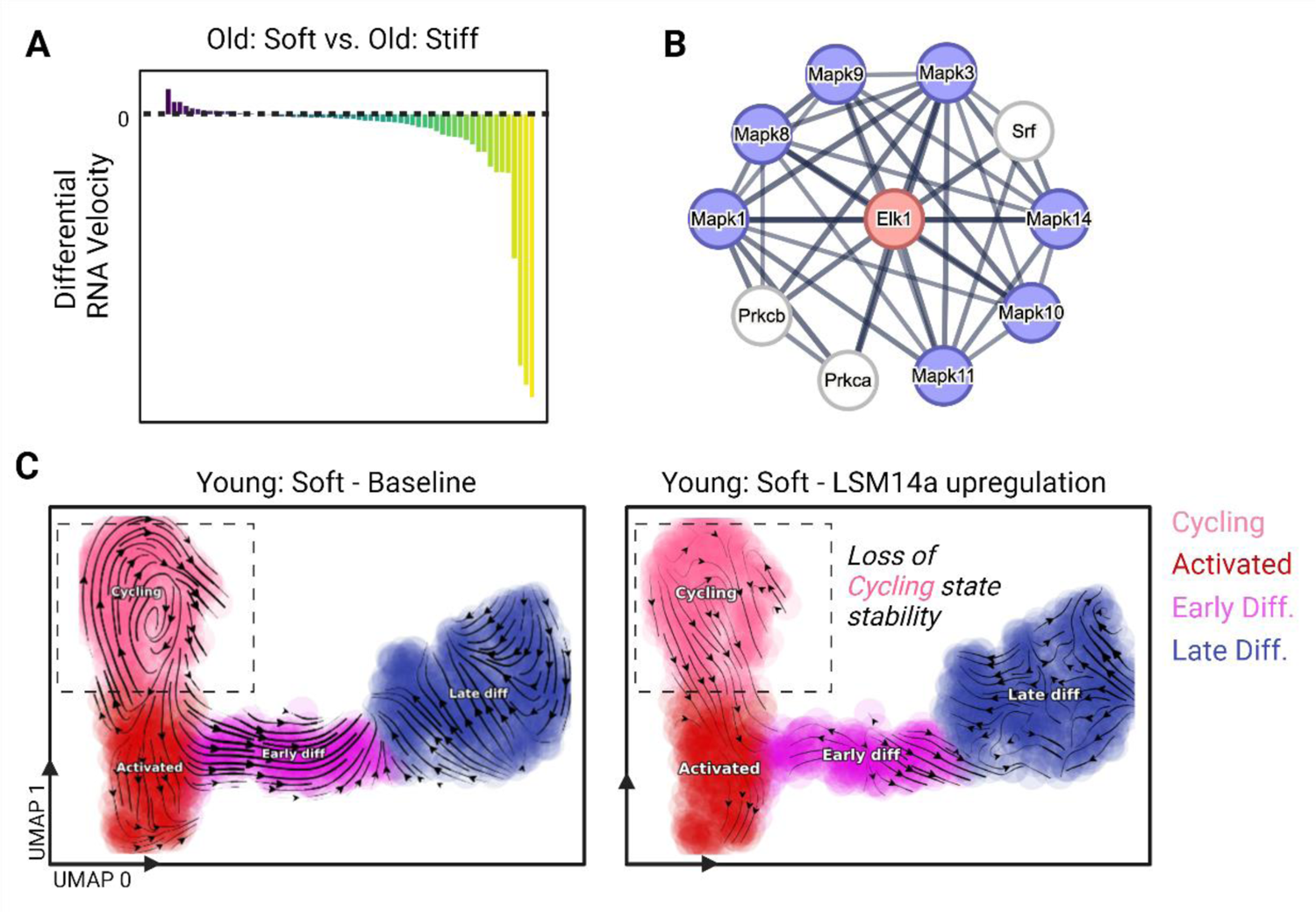
(**A**) Random forest identification of differential RNA velocities between old MuSCs on soft versus stiff substrates. (**B**) Protein interaction network of ETS Like-1 protein Elk-1 (ELK1) using STRING. The confidence of interaction was set to high when determining ELK1 interactions (>0.9 out of 1.0). (**D**) *In silico* perturbation of vector fields projected onto UMAP of upregulated *LSM14a* in young MuSCs on soft matrices. Cluster colors correspond to stage of lineage progression. Only significant vectors are shown.

## Supplemental File Legends

**Supplemental File 1**. Gene sets from subclusters identified from aggregate UMAP clustering.

## STAR Methods

All reagents, equipment, and software code used are outlined in the **Key Resources Table** and will be available upon publication.

### Resource Availability

#### Lead Contact

Further information and requests for resources should be directed to and will be fulfilled by the lead contact, Fabrisia Ambrosio.

#### Materials availability

There were no new unique reagents used in this study.

#### Data and code availability

All data used to support the conclusions of this study are available from the corresponding author upon reasonable request. This paper does not report original code. Information regarding Monocle 3.0 (Monocle3 (RRID:SCR_018685), RNA velocity (RRID:SCR_018167), and *dynamo* (RRID:SCR_017541), can be found using the SciCrunch Registry. The scRNA-seq data that support the findings of this study will be available upon publication.

### Experimental model and subject details

#### Animals

C57BL/6J young (4-6 months old) and aged (22-24 months old) male mice were received from either the NIA rodent colony or Jackson Laboratories. All animal studies were approved by the Institutional Animal Care and Use Committee of the University of Pittsburgh. Mice were housed in pathogen-free cages with a temperature and humidity-controlled room with a 12-hour light/dark cycle. Each cage received enrichment items, and food and water were provided *ad libitum*. On average, 3-5 mice were housed per cage. The animals were assessed before the studies, and any animals with obvious signs of health problems (e.g., lethargy, poor grooming, 20% weight loss, or hunching) were excluded. All animals were then randomly assigned to intervention groups and ear tagged. All investigators performing endpoint analyses were blinded to the experimental group whenever possible.

#### BAPN administration and cardiotoxin injuries

Sterile saline or BAPN (290 μg/μl in sterile saline; approximately 550 μg/g of body weight; CDS007521; Sigma, Burlington, MA, USA) was subcutaneously injected daily into the nape of the neck of aged mice for a total of six weeks. Young mice received sterile saline injections alone for the same timeframe to determine how aged BAPN injected mice compared to young. Four weeks following the start of BAPN and/or saline injections, TA muscles were injured bilaterally via intramuscular injection of cardiotoxin (10 μg/ml; 217503; Sigma). Two weeks following injury, TA muscles were assessed for force producing capacity using in-situ contractile testing, and then were immediately harvested, frozen, and stored at −80°C until further analysis.

#### In situ contractile testing

Contractile testing was performed 14 days after injury using an *in-situ* testing apparatus (Aurora (Model 809B, Aurora Scientific Inc, Canada), stimulator (Model 701C, Aurora Scientific Inc), and force transducer (Aurora Scientific Inc). Animals were anesthetized using 1.5-3% isoflurane prior to the functional testing. The right Achilles tendon was cut, and the peroneal nerve was exposed via a small incision on the right knee. Mice were then placed supine on a platform with the foot to be tested positioned at 20° of plantarflexion and affixed to the footplate using cloth tape. The foot was stimulated six times at a single frequency (60Hz) by placing a hook electrode on the peroneal nerve. Half-relaxation time was quantified using the single twitch data. Next, successive tetanic contractions at 10, 30, 50, 80, 100, 120, 150, 180, and 200 Hz were elicited to obtain a force-frequency curve, with a 2-minute rest between contractions, as per our previous reports (90, 95). Peak specific tetanic force was quantified by extracting the highest force value from the series of tetanic contractions of each mouse. Animals displaying evidence of external injuries or tumor growths were excluded from analysis. Functional testing was repeated by two independent investigators over three cohorts of animals to ensure reproducibility.

## Method Details

### Biaxial Testing

Biaxial tensile testing of the gastrocnemius muscle was conducted as previously described (8). Briefly, the medial head of the gastrocnemius muscle was collected from young and aged mice and immediately placed in Ringer’s solution for at least one hour before mechanical testing. Muscle samples were cut into squares (5 mm x 5 mm) after measuring the sample thickness at five individual points. Samples were then attached to a 50 g load cell with two loops of suture connected to each side, with four hooks per side (see Fig. 1A). The sample deformation was measured by real-time tracing of coordinates from a four-marker array positioned in the center of each muscle sample. Gradient tensor F was calculated from the shape functions as previously described(18). An equi-stress protocol was executed with the specimens submerged in PBS at room temperature utilizing a tare load of 0.5 gram and a cycle time of 10 seconds each. Samples were then tested up to the determined stress of 32 kPa, which was found to be the upper limit muscles can tolerate without permanent deformation. All data were referenced to the pre-conditioned float state.

#### In-plane green’s strain energy map

A structural determinist approach was selected to quantify single fiber Green’s strain (E) within the gastrocnemius muscles tested under biaxial tensile states (96). First, fiber network models were generated as stochastically equivalent to experimental samples based on the fibril network architectural features previously evaluated using simple harmonic generation imaging (8). Fiber intersection density, diameter, and orientation index were the input parameters of a mesh generator. Artificial fiber networks were then matched with the distribution of fiber nodes, orientation, and connectivity, and then extracted with digital image processing (18, 97). Taking advantage of the biaxial test symmetry, square fiber network meshes were measured 500 µm by 500 µm to reduce the cost of the computational analysis (20). Biaxial tensile tests were then simulated in Abaqus 6.14 (Dassault Systems, Waltham, MA), as described (20). Briefly, symmetry boundary conditions were set on two edges of the square mesh, where axial displacements were 0, while axial loads from biaxial tests were applied on the remaining edges. All fibers were computationally modelled with straight B22 beam elements, and the Yeoh hyper-elastic material model was assigned (98). At the end of each analysis, results were automatically extracted with a custom Python module to calibrate the Yeoh material parameters by fitting the experimental biaxial data (98). After calibrating material properties, finite element analyses were run to evaluate E_f_ maps and histograms under the biaxial tensile tests.

#### Hydroxyproline assay

The concentration of mature crosslinked and immature crosslinked and non-crosslinked collagen was determined using previously established protocols (99). Briefly, freshly harvested TA muscles from saline and/or BAPN treated mice were weighed, snap-frozen, and pulverized with a mortar and pestle over dry ice. The samples were then rinsed in 1 mL of PBS with agitation for 30 minutes at 4°C, and then centrifuged at 16,000g for 30 minutes at 4°C. Following centrifugation, PBS was replaced with a 1:6 (weight: volume) solution of 0.5 M acetic acid (984303; Thermo Fisher, Waltham, MA, USA) with 1 mg/ml pepsin (20343; Thermo Fisher) and stirred overnight at 4°C. The following day, the samples were then centrifuged at 16,000g for 30 min at 4°C. Next, the supernatant was collected as the pepsin-soluble fraction (PSF), or non-cross-linked and immature crosslinked collagen, and the pellet was kept as the pepsin-insoluble fraction (PIF), or mature crosslinked collagen. A 1:1 volume of 4 M NaCl was added to the PSF and incubated with agitation for 30 minutes at 4°C before 30 minutes of centrifugation at 16,000g at 4°C. The supernatant was then discarded, and the collagen fractions were measured using a Hydroxyproline Assay Kit (ab222941; Abcam, Cambridge, UK) per the manufacturer’s instructions.

#### Analysis of muscle regeneration

Muscle tissues were frozen by completely immersing in liquid nitrogen-cooled 2-Methylbutane for 1 minute. Frozen muscles were sectioned at 10 µm thickness using CryoStar NX50 Cryostat (Thermo Fisher,). Muscle sections were then fixed in 2% paraformaldehyde solution (J19943K2, Thermo Fisher) diluted in PBS followed by three 5-minute PBS washes at room temperature. Samples were permeabilized with 0.01% Triton X-100 in PBS for 15 minutes and blocked with 3% bovine serum albumin (A7906, Sigma) in 0.01% Triton X-100 in PBS (blocking buffer) for 1 hour at room temperature. The sample slides were incubated with rat anti-Laminin (ab79057, Abcam, Cambridge, UK) in 5% goat serum (191356, MP Biomedicals, Solon, OH) in blocking buffer overnight at 4 °C. The slides were washed for 5 minutes 3 times with PBS followed by incubation with goat anti-Rat IgG (H+L) Alexa Fluor 594 secondary antibody (A11012, Thermo Fisher) at 1:500 dilution for one hour at room temperature. After three 5-minute PBS washes, the samples were incubated with 4’,6-diamidino-2-phenylindole (DAPI) (422801, BioLegend, San Diego, CA) for 2 minutes at room temperature. The slides were washed once with PBS for 5 minutes before coverslips (12-545-100, Thermo Fisher) were mounted with Gelvatol mounting media (81365; Sigma) and let to dry. Images were taken using inverted microscopy (Observer Z1, Carl Zeiss AG, Oberkochen, Germany) with a 20x objective lens. The area of centrally nucleated myofibers was measured using ImageJ software using a previously established protocol (100).

#### Primary muscle stem cell (MuSC) isolation and culture

MuSCs were freshly isolated from young/aged mice for each experiment using previously established protocols (101). Briefly, forelimb and hindlimb musculature were dissected and placed in ice cold wash media (Hanks Balanced Saline Solution (HBSS); Thermo Fisher, 10% Fetal Bovine Serum (FBS); Thermo Fisher, 1% Penicillin-Streptomycin; Thermo Fisher). Muscles were then finely chopped and then incubated with agitation in 750 U/mL of Collagenase II (Gibco) for one hour, followed by an additional 30 minutes in 1000 U/mL Collagenase II and 100 U/mL Dispase (Gibco). Mononuclear cells were then collected through a 40 μm strainer (Thermo Fisher), and immunolabeled with antibodies for CD31, CD45, Sca1, and Integrin α7, as previously described (11-0311-82; 11-0451-82; 25-5981-82; MA5-23555, respectively; Thermo Fisher) (101). Propidium iodide was used as a dead cell marker (1mg/ml aqueous solution; 25535-16-4, Alfa Aesar, Lancashire, UK). Cells were then sorted using fluorescence-activated cell sorting (BD Biosciences; Franklin Lakes, New Jersey, USA) based on live/dead and negative expression of CD31, CD45, and Sca1.

#### Quantification of nuclear morphology

For the quantification of nuclear morphology, DAPI-labeled cross sections and isolated MuSCs were obtained at 40x magnification using an A1 confocal microscope (Nikon, Tokyo, Japan). Afterwards, image processing and morphometric feature extraction were performed using CellProfiler™ software (v4.0, The Broad Institute). Fifty-three shape features of nuclei were determined using the “identify primary objects” followed by the “measure object size shape” and “export to spread sheet” module. Principal Component Analysis (PCA; unsupervised) and PCA-linear discriminant analysis (PCA-LDA; supervised) were performed to facilitate the identification of segregation of the nuclear morphological features identified by CellProfiler™ software (23). In PCA-LDA analysis, the small number of PCs corresponds to sample size in each group, were included to prevent overfitting. To determine variables of nuclear shape contributing to PCs, loading matrix, a correlation between the original variable and PCs, were extracted.

#### PDMS fabrication

PDMS substrates with different stiffnesses were prepared as previously described. Briefly, Sylgard 527 was prepared by mixing 1) equal weights or 2) 1:1.8 weight ratio of Dielectric Gel Part A and Part B and mixed well manually for 12 kPa and 29 kPa substrates, respectively (NC1208196; Thermo Fisher). The mixtures were degassed using a vacuum desiccator for 30 minutes. Next, PDMS was poured into 60 mm diameter Petri dishes to make ∼1.2 mm thick substrate for cell culture. Alternatively, 30 µL of PDMS was poured onto an 18 mm circular coverslip followed by a one-minute spin at 930 rpm using a spin coater (Laurell WS-650SZ-6NPP/A1/AR1) to make ∼ 100 µm-thick films for cell culture for immunocytochemistry imaging of cell on PDMS substrates. PDMS substrates were cured at 60 °C for 16 hours for all experiments.

PDMS substrates were treated with plasma cleaner (Harrick Plasma, Ithaca, NY) at the medium power setting for 20 seconds and soaked in water until the next step. Water was replaced with 70% ethanol and sterilized for 30 minutes under UV light in a cell culture hood. PDMS substrates were rinsed with sterile water three times and coated with 50 µg/mL of fibronectin (F4759, Sigma-Aldrich, St. Louis, MO) diluted in water for one hour at room temperature. Fibronectin solution was then removed, and the substrates were rinsed with water twice, followed by a PBS rinse before cell seeding. PDMS substrates were never left dry after plasma treatment. Cells were seeded on PDMS substrates at ∼ 160 – 200 cells/mm^2^.

#### Immunofluorescence and image acquisition

For the nuclear morphology image analysis, freshly isolated young MuSCs were cultured on soft or stiff PDMS substrates for a week before nuclear staining and imaging of the cells on the PDMS substrates to avoid potential alterations in cell morphology by transfer of the cells. For the staining of other experiments for morphological analysis of adherent cells, MuSCs were fixed and stained with DAPI two days after seeding, which is just enough time to allow MuSC adhesion to the substrate. The following antibodies were used for the other immunostaining studies: Mouse anti-PAX7 antibody at 5 µg/mL (DSHB, Iowa, U.S.A.) and Rabbit anti-MyoD antibody at 1:500 (sc-760, Santa Cruz, Texas, U.S.A.).

#### Imaging flow cytometry and analysis

Isolated MuSCs were plated on soft/stiff substrates and cultured for seven days. Next, the cells were trypsinized with 0.25% trypsin for one minute at 37°C and collected in 1.5 mL Eppendorf tubes for subsequent analysis. The pelleted cells were fixed, permeabilized, blocked, and immunolabeled for myogenic markers of Pax7 and MyoD. Labeled cells were imaged at single-cell resolution using imaging flow cytometry (ISX; Flow cytometry core, Department of Immunology, University of Pittsburgh). ISX was conducted using a 60X objective at a resolution of 0.3 μm^2^/pixel. Filtered sheath buffers were used to ensure the absence of debris and non-cellular components. Samples were acquired using INSPIRE® software with the highest sensitivity by acquiring images at the lowest speed. All lasers employed were used at the optimal power settings based on an unstained cell control. Gating was performed on unstained control cells for acquiring images of single and focused cells. For analyzing myogenic markers, IDEAS software was used by performing a nuclear co-localization step to detect the true positives (i.e., Pax7 and MyoD fluorescent intensity) in the imaging dataset. Nuclear morphology features were extracted using IDEAS software per cell, which were used for downstream computational analysis. Every experiment was performed in at least two independent runs. In every run, three sets of imaging data were acquired. Data were presented as an average of the three sets of imaging data in one representative run.

#### Single cell RNA sequencing

Both young and old MuSCs were collected at the same timepoints after culturing on soft or stiff surfaces for single cell RNA sequencing (scRNA-seq) to minimize batch effects (102). A total of three mice per age group were used for cell isolations to ensure biological diversity, and a total of three technical replicates per substrate condition were used to mitigate experimental error. Following the collection of cells from PDMS experiments, MuSCs were loaded into a 10X Chromium Controller using the Single Cell 3’ Reagent kit v3 per manufacturer’s instructions (10x Genomics, Pleasanton, CA, USA). Resulting libraries were sequenced on an Illumina NovaSeq 6000 platform yielding a read depth of 200 million reads/sample (Novogene, China).

#### Single cell RNA sequencing data preprocessing

The Cell Ranger pipeline (6.1.2) was utilized to align each sample’s reads to the mouse reference genome (mm10). The unspliced and spliced counts for each sample were obtained using *velocyto* (0.17.17). For downstream analysis, the gene filtering and normalization was performed using *dynamo* (1.1.0, https://github.com/aristoteleo/dynamo-release) in python. For each sample, the cells filtered out with the Monocle 3.0 preprocessing were also filtered out for *dynamo* downstream analysis. Briefly, the filtration of gene counts and cells were based on the following criteria: genes expressed in only 1.0% of total cells, cells expressing >20.0% of mitochondrial-related gene counts, cells expressing <5% of ribosomal-related reads, and trace amounts of non-myogenic cells (i.e., fibroadipo-progenitor cells (*Pdgfra*), macrophages (*Adgre1*) tenocytes (*Tnmd*), and neuronal cells (*Mmp17*), For downstream analysis a total of 5,913 cells and 32,285 genes were obtained from all samples. In the *dynamo* preprocessing step (recipe monocle), the top 4000 genes were utilized for RNA velocity calculation following size normalization and log1p transformation. For visualization purposes, dimensionality reduction was performed with UMAP on the 30-dimensional PCA space that was obtained from *dynamo*’s *reduceDimension* function.

#### Dimensionality Reduction

For gene expression, combining all cells from each sample and selecting the intersection of all genes that were labeled as “use_for_dynamics” in *dynamo* results in a 5,913 cell by 1363 gene expression matrix. With this, UMAP was performed. For RNA velocity, combining all cells from each sample and selecting the intersection of all genes that were labeled as “use_for_dynamics” in *dynamo* results in a 5,913 cell by 1363 gene expression matrix. Extracting the RNA velocity for each gene for each cell results in a 5,913 cell by 1363 gene velocity matrix. With this, UMAP was performed.

#### Classifying cells

The classification of cells into the one of the four groups, cycling, activated, early differentiation, and late differentiation was obtained using a combination of unsupervised graph-based clustering as well as manual characterization using previously reported state-specific markers (2, 56, 103). Following dimensionality reduction using PCA and visualization using UMAP, the Louvain clustering algorithm was performed to generate clusters based on discrete cluster-specific gene expression. The corresponding gene sets were then used to 1) corroborate cluster annotation with previously published datasets and 2) to generate bubble plots indicating cluster-specific gene expression. Bubble plots were generated following cluster classification by determining the average expression of each gene of interest (GOI) per cluster, followed by measuring the percentage of cells within each cluster expressing each GOI.

#### Monocle 3.0

Trajectory analysis was performed using Monocle 3.0 using the same preprocessed data as used for *dynamo* analyses. Briefly, following preprocessing (see Preprocessing MuSC scRNA-seq data), dimensionality reduction was performed using UMAP. To confirm annotation of MuSC subclusters using Monocle 3.0 was the same as conventional dimensionality reduction approaches, the trajectory was probed for expression of genes corresponding to each cellular state (i.e., activated, cycling, early differentiation, and late differentiation). Following annotation, cells with the highest expression of *Pax7* were selected as the start point for pseudotime progression. Cells with high expression of Pax7 *in vivo* have been shown to represent MuSCs with the highest degree of pluripotency (104). Pseudotime progression in the aggregated dataset (i.e., inclusion of all four groups) corroborated our subcluster annotation by showing both self-renewing and differentiating trajectories. A correlation analysis of gene expression along pseudotime progression was performed to identify genes expressed throughout each trajectory, which are shown in Fig. 3A, as well as the bubble plot in Fig. 3B. Following aggregate trajectory characterization, individual trajectory analysis for each sample was performed.

#### Differential analysis

The gene specific analysis (GSA) feature of Partek Flow was used for differential analysis of gene expression between samples, which is equivalent to limma-trend (102) (Partek Genomics, St. Louis, MO, USA). The resulting gene sets were filtered using an FDR step up (p.adj) of 0.05. Genes +/- 2-fold change were determined to be statistically significant.

#### RNA velocity

RNA velocity estimations were made using *dynamo*, as previously described (41). Briefly, reads were aligned to the mm10 mouse reference transcriptome with *cellranger,* and *velocyto* was utilized to quantify the read counts of both pre-mRNA (unspliced; mapped exon-intron boundaries, or the intron of the transcript alone) and mature mRNA (spliced; mapping only to exonic regions of transcripts). To remain consistent with the previous analysis performed with Monocle 3.0, cells filtered out in the Monocle 3.0 analysis were also filtered out for the *dynamo* analyses. The following set of ordinary differential equations model the turnover dynamics of a specific gene *i* that is actively transcribed in a cell,

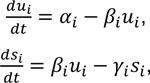

where *u_i_* represents the number of unspliced RNA copies of gene *i*, 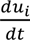 is the rate of production of unsliced transcripts, *s_i_* represents the number of spliced RNA copies, 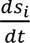 is the rate at which spliced RNA copies are produced (i.e., RNA velocity), *α_i_* is the rate of transcription, *β_i_* is the rate constant of transcript splicing, and *γ_i_* is the rate constant of transcript degradation. *u_i_* and *s_i_* were both measured in scRNA-seq. Furthermore, with 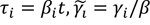, one can define a scaled RNA velocity of a gene relative to its own splicing time scale, 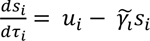. The scaled gene specific degradation rate constant γ_i_ was determined using Ordinary Least Squares regression as in the original RNA velocity paper of La Manno et al. (40), in which a pseudo steady state was assumed to lie in the *u*-*s* phase space where cells have extreme high unspliced and high spliced RNA expression.

#### Continuous vector field reconstruction

Detailed methodology on vector field reconstruction has been published previously by Qui et al (41). In brief, *dynamo* aims to reconstruct a continuous vector field from discrete gene expression data obtained from scRNA-seq data and instantaneous RNA velocities. Notice that *α* is a function of the concentrations of the elements regulating a gene under consideration, so a function of ***s*** under a Markovian assumption (105), and 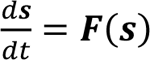, where the vector **s** represents the gene expression, 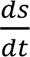 represents the RNA velocity, and **F** represents the learned vector field function, containing genome wide regulation, mapping any point in the gene expression space to the RNA velocity space.

#### Vector field differential geometry analysis

We projected the reconstructed continuous vector field onto a two-dimensional UMAP space, then identified fixed points that indicate the behavior and direction a cell at a specific cellular state tends to move along the transcriptional landscape. In other words, one can predict on the transition between cell states over an arbitrary time scale.

As before, the details of these calculations and derivations are available in the original *dynamo* publication (41). In brief, a fixed point is where the value of the vector field function is zero. The eigenvalues of the Jacobian of that point was evaluated to categorized that point as either a stable fixed point (attractor), unstable fixed point (repulsor), or a saddle point. From the Jacobian obtained from the vector field, one can characterize fixed points and observe where cells end up in the cell state space. With this, we can then ask the question “(1) did the cell reach a point in space where results in no changes in the cell state (i.e., stable), or (2) did the cell continue to change states in space indefinitely (i.e., unstable)?”

#### Random forests classification

Given the emergence of stable points in the cycling and activated cell states for old MuSCs on a soft substrate compared to stiff, each of these states were selected for differential analyses using random forest classification. To reduce the number of features, the intersection of all genes that were labeled as “use_for_dynamics” in *dynamo* were selected for a total of 1363 genes. With this set of cells and genes, the corresponding RNA velocities for each gene per each cell were utilized as the input data for the random forests. Ten rounds of random forests, using *sklearn*, were performed (n_estimators = 100, max_features = ‘sqrt’, max_depth = 1, class_weight = ‘balanced’) with a 0.8 and 0.2 training and testing data split, respectively. The max depth of 1 was chosen to extract the genes that displayed the greatest degree of differential velocities between groups. For each round of random forests, the important features were extracted.

#### Gene set enrichment and TRANSFAC analysis

Following gene set generation from either random forest classification or unique biomarkers per subcluster classification, genes were imported into gProfiler (https://biit.cs.ut.ee/gprofiler/) in rank-order. Prior to analysis, multiple testing was corrected for using a Benjamini-Hochberg false discovery rate of >0.05. To account for overlap in synonymous biological processes, a term size limit greater than 5 and less than 500 was selected (106). To identify predicted transcription factors common to nucleotide sequences of the inputted gene sets, TRANSFAC was used with similar testing correction parameters (47).

#### Protein-protein interaction networks

STRING (https://string-db.org/) was used to infer physical binding partners of LSM14a, as well as signaling partners of Elk1, as has been previously described. Briefly, each gene of interest (GOI) was input into the STRING database, and settings were made according to each GOI. For LSM14a, “physical subnetwork” and “high confidence” of interaction scores was selected, along with a maximum of no more than 20 interactions. For Elk1, “full STRING network”, “high confidence” of interactions, and no more than 10 interactions were selected for network generation.

#### Vector field in silico perturbations

The Jacobian can also be utilized to understand gene-gene regulations such as whether a gene upregulates or down regulates another gene. Mathematically, a Jacobian element 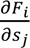 encodes how the transcriptional velocity of gene i is impacted by changes in expression of gene j. Consider that the expression state of a cell changes form **s** to **s** + Δ**s** through, e.g., adding one or some of the transcripts ectopically by amount Δ**s**, then the vector field changes approximately to,

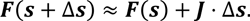

Thus **J** ⋅ Δ**s** propagates the genetic perturbation Δ**s** through the gene regulatory network.

### Quantification and statistical analysis

#### Statistics

Sample size determination for CellProfiler, ImageStream, and *in vivo* analyses were determined from our previously published studies (29, 90, 95). For experiments using isolated MuSCs requiring a power level of 0.08, n=3 mice/age were used. For *in vivo* experiments requiring an alpha level of 0.08, n=5/mice/age were used. For scRNA-seq experiments, isolated MuSCs from n=3 mice/age were used for *in vitro* experiments, which were split into three technical replicates prior to sequencing. A read depth of 200 million reads/sample was collected to ensure adequate power (107). Statistical analyses were performed using GraphPad Prism (v. 9.0). Significance was assumed at an alpha level of 0.05. Kolmogorov-Smirnov test and F-test were initially performed to assess normality and equality of variance, respectively. For all ANOVA analyses resulting in a p-value of less than 0.05, appropriate post-hoc tests were performed and described in figure legends. For multiple comparisons, Bonferroni correction methods were performed to mitigate type I error inflation.

## References

1. Blau HM, Cosgrove BD, and Ho AT. The central role of muscle stem cells in regenerative failure with aging. Nat Med. 2015;21(8):854–62.

2. Kimmel JC, Hwang AB, Scaramozza A, Marshall WF, and Brack AS. Aging induces aberrant state transition kinetics in murine muscle stem cells. Development. 2020;147(9).

3. Wang J, Zhang K, Xu L, and Wang E. Quantifying the Waddington landscape and biological paths for development and differentiation. Proceedings of the National Academy of Sciences. 2011;108(20):8257–62.

4. Goldberg AD, Allis CD, and Bernstein E. Epigenetics: a landscape takes shape. Cell. 2007;128(4):635–8.

5. Engler AJ, Sen S, Sweeney HL, and Discher DE. Matrix elasticity directs stem cell lineage specification. Cell. 2006;126(4):677–89.

6. Gilbert PM, Havenstrite KL, Magnusson KE, Sacco A, Leonardi NA, Kraft P, et al. Substrate elasticity regulates skeletal muscle stem cell self-renewal in culture. Science. 2010;329(5995):1078-81.

7. Gosselin LE, Adams C, Cotter TA, McCormick RJ, and Thomas DP. Effect of exercise training on passive stiffness in locomotor skeletal muscle: role of extracellular matrix. J Appl Physiol (1985). 1998;85(3):1011-6.

8. Stearns-Reider KM, D’Amore A, Beezhold K, Rothrauff B, Cavalli L, Wagner WR, et al. Aging of the skeletal muscle extracellular matrix drives a stem cell fibrogenic conversion. Aging Cell. 2017;16(3):518–28.

9. Zimmerman SD, McCormick RJ, Vadlamudi RK, and Thomas DP. Age and training alter collagen characteristics in fast- and slow-twitch rat limb muscle. J Appl Physiol (1985). 1993;75(4):1670-4.

10. Freitas-Rodríguez S, Folgueras AR, and López-Otín C. The role of matrix metalloproteinases in aging: Tissue remodeling and beyond. Biochim Biophys Acta Mol Cell Res. 2017;1864(11 Pt A):2015-25.

11. Lacraz G, Rouleau AJ, Couture V, Söllrald T, Drouin G, Veillette N, et al. Increased Stiffness in Aged Skeletal Muscle Impairs Muscle Progenitor Cell Proliferative Activity. PLoS One. 2015;10(8):e0136217.

12. Wood LK, Kayupov E, Gumucio JP, Mendias CL, Claflin DR, and Brooks SV. Intrinsic stiffness of extracellular matrix increases with age in skeletal muscles of mice. J Appl Physiol (1985). 2014;117(4):363-9.

13. Mann CJ, Perdiguero E, Kharraz Y, Aguilar S, Pessina P, Serrano AL, et al. Aberrant repair and fibrosis development in skeletal muscle. Skeletal Muscle. 2011;1:21-.

14. Rosant C, Nagel MD, and Pérot C. Aging affects passive stiffness and spindle function of the rat soleus muscle. Exp Gerontol. 2007;42(4):301–8.

15. Gao Y, Kostrominova TY, Faulkner JA, and Wineman AS. Age-related changes in the mechanical properties of the epimysium in skeletal muscles of rats. J Biomech. 2008;41(2):465–9.

16. Schüler SC, Kirkpatrick JM, Schmidt M, Santinha D, Koch P, Di Sanzo S, et al. Extensive remodeling of the extracellular matrix during aging contributes to age-dependent impairments of muscle stem cell functionality. Cell Rep. 2021;35(10):109223.

17. Meyer GA, and Lieber RL. Elucidation of extracellular matrix mechanics from muscle fibers and fiber bundles. Journal of biomechanics. 2011;44(4):771–3.

18. D’Amore A, Amoroso N, Gottardi R, Hobson C, Carruthers C, Watkins S, et al. From single fiber to macro-level mechanics: A structural finite-element model for elastomeric fibrous biomaterials. J Mech Behav Biomed Mater. 2014;39:146–61.

19. Carleton JB, D’Amore A, Feaver KR, Rodin GJ, and Sacks MS. Geometric characterization and simulation of planar layered elastomeric fibrous biomaterials. Acta Biomater. 2015;12:93–101.

20. D’Amore A, Nasello G, Luketich SK, Denisenko D, Jacobs DL, Hoff R, et al. Meso-scale topological cues influence extracellular matrix production in a large deformation, elastomeric scaffold model. Soft Matter. 2018;14(42):8483–95.

21. Pajerowski JD, Dahl KN, Zhong FL, Sammak PJ, and Discher DE. Physical plasticity of the nucleus in stem cell differentiation. Proc Natl Acad Sci U S A. 2007;104(40):15619–24.

22. Spagnol ST, and Dahl KN. Active cytoskeletal force and chromatin condensation independently modulate intranuclear network fluctuations. Integr Biol (Camb). 2014;6(5):523–31.

23. McQuin C, Goodman A, Chernyshev V, Kamentsky L, Cimini BA, Karhohs KW, et al. CellProfiler 3.0: Next-generation image processing for biology. PLoS Biol. 2018;16(7):e2005970.

24. Heckenbach I, Mkrtchyan GV, Ezra MB, Bakula D, Madsen JS, Nielsen MH, et al. Nuclear morphology is a deep learning biomarker of cellular senescence.

25. Meschiari CA, Ero OK, Pan H, Finkel T, and Lindsey ML. The impact of aging on cardiac extracellular matrix. Geroscience. 2017;39(1):7–18.

26. Willett TL, Dapaah DY, Uppuganti S, Granke M, and Nyman JS. Bone collagen network integrity and transverse fracture toughness of human cortical bone. Bone. 2019;120:187–93.

27. Aicher BO, Zhang J, Muratoglu SC, Galisteo R, Arai AL, Gray VL, et al. Moderate aerobic exercise prevents matrix degradation and death in a mouse model of aortic dissection and aneurysm. Am J Physiol Heart Circ Physiol. 2021;320(5):H1786–h801.

28. Spengler DM, Baylink DJ, and Rosenquist JB. Effect of beta-aminopropionitrile on bone mechanical properties. J Bone Joint Surg Am. 1977;59(5):670–2.

29. Iijima H, Gilmer G, Wang K, Bean AC, He Y, Lin H, et al. Age-related matrix stiffening epigenetically regulates α-Klotho expression and compromises chondrocyte integrity. Nat Commun. 2023;14(1):18.

30. Trapnell C, Cacchiarelli D, Grimsby J, Pokharel P, Li S, Morse M, et al. The dynamics and regulators of cell fate decisions are revealed by pseudotemporal ordering of single cells. Nat Biotechnol. 2014;32(4):381–6.

31. Qiu X, Hill A, Packer J, Lin D, Ma YA, and Trapnell C. Single-cell mRNA quantification and differential analysis with Census. Nat Methods. 2017;14(3):309–15.

32. Bentzinger CF, Wang YX, Maltzahn J, Soleimani VD, Yin H, and Rudnicki M. Fibronectin regulates Wnt7a signaling and satellite cell expansion. Cell Stem Cell. 2013;12.

33. Baldin V, Lukas J, Marcote MJ, Pagano M, and Draetta G. Cyclin D1 is a nuclear protein required for cell cycle progression in G1. Genes Dev. 1993;7(5):812–21.

34. Charge SB, and Rudnicki MA. Cellular and molecular regulation of muscle regeneration. Physiol Rev. 2004;84(1):209–38.

35. Núñez-Álvarez Y, Hurtado E, Muñoz M, García-Tuñon I, Rech GE, Pluvinet R, et al. Loss of HDAC11 accelerates skeletal muscle regeneration in mice. Febs j. 2021;288(4):1201–23.

36. Edfors F, Danielsson F, Hallström BM, Käll L, Lundberg E, Pontén F, et al. Gene-specific correlation of RNA and protein levels in human cells and tissues. Mol Syst Biol. 2016;12(10):883.

37. Gry M, Rimini R, Strömberg S, Asplund A, Pontén F, Uhlén M, et al. Correlations between RNA and protein expression profiles in 23 human cell lines. BMC Genomics. 2009;10(1):365.

38. Liu Y, Beyer A, and Aebersold R. On the Dependency of Cellular Protein Levels on mRNA Abundance. Cell. 2016;165(3):535–50.

39. Bevilacqua A, Ceriani MC, Capaccioli S, and Nicolin A. Post-transcriptional regulation of gene expression by degradation of messenger RNAs. J Cell Physiol. 2003;195(3):356–72.

40. La Manno G, Soldatov R, Zeisel A, Braun E, Hochgerner H, Petukhov V, et al. RNA velocity of single cells. Nature. 2018;560(7719):494-8.

41. Qiu X, Zhang Y, Martin-Rufino JD, Weng C, Hosseinzadeh S, Yang D, et al. Mapping transcriptomic vector fields of single cells. Cell. 2022;185(4):690–711.e45.

42. Díaz-Uriarte R, and Alvarez de Andrés S. Gene selection and classification of microarray data using random forest. BMC Bioinformatics. 2006;7(1):3.

43. Wek RC. Role of eIF2α Kinases in Translational Control and Adaptation to Cellular Stress. Cold Spring Harb Perspect Biol. 2018;10(7).

44. Tikhonova EB, Gutierrez Guarnizo SA, Kellogg MK, Karamyshev A, Dozmorov IM, Karamysheva ZN, et al. Defective Human SRP Induces Protein Quality Control and Triggers Stress Response. J Mol Biol. 2022;434(22):167832.

45. Gille H, Strahl T, and Shaw PE. Activation of ternary complex factor Elk-1 by stress-activated protein kinases. Curr Biol. 1995;5(10):1191–200.

46. Bresson S, Shchepachev V, Spanos C, Turowski TW, Rappsilber J, and Tollervey D. Stress-Induced Translation Inhibition through Rapid Displacement of Scanning Initiation Factors. Mol Cell. 2020;80(3):470–84.e8.

47. Matys V, Kel-Margoulis OV, Fricke E, Liebich I, Land S, Barre-Dirrie A, et al. TRANSFAC and its module TRANSCompel: transcriptional gene regulation in eukaryotes. Nucleic Acids Res. 2006;34(Database issue):D108-10.

48. Bernet JD, Doles JD, Hall JK, Kelly Tanaka K, Carter TA, and Olwin BB. p38 MAPK signaling underlies a cell-autonomous loss of stem cell self-renewal in skeletal muscle of aged mice. Nat Med. 2014;20(3):265–71.

49. Durand S, Cougot N, Mahuteau-Betzer F, Nguyen CH, Grierson DS, Bertrand E, et al. Inhibition of nonsense-mediated mRNA decay (NMD) by a new chemical molecule reveals the dynamic of NMD factors in P-bodies. J Cell Biol. 2007;178(7):1145–60.

50. Hug N, Longman D, and Cáceres JF. Mechanism and regulation of the nonsense-mediated decay pathway. Nucleic Acids Res. 2016;44(4):1483–95.

51. Wallmeroth D, Lackmann J-W, Kueckelmann S, Altmüller J, Dieterich C, Boehm V, et al. Human UPF3A and UPF3B enable fault-tolerant activation of nonsense-mediated mRNA decay. The EMBO Journal. 2022;41(10):e109191.

52. Hu G, Cui K, Northrup D, Liu C, Wang C, Tang Q, et al. H2A.Z facilitates access of active and repressive complexes to chromatin in embryonic stem cell self-renewal and differentiation. Cell Stem Cell. 2013;12(2):180–92.

53. Huh YH, Noh M, Burden FR, Chen JC, Winkler DA, and Sherley JL. Sparse feature selection identifies H2A.Z as a novel, pattern-specific biomarker for asymmetrically self-renewing distributed stem cells. Stem Cell Res. 2015;14(2):144–54.

54. Tanaka KK, Hall JK, Troy AA, Cornelison DD, Majka SM, and Olwin BB. Syndecan-4-expressing muscle progenitor cells in the SP engraft as satellite cells during muscle regeneration. Cell Stem Cell. 2009;4(3):217–25.

55. Sousa-Victor P, Gutarra S, Garcia-Prat L, Rodriguez-Ubreva J, Ortet L, Ruiz-Bonilla V, et al. Geriatric muscle stem cells switch reversible quiescence into senescence. Nature. 2014;506(7488):316-21.

56. Oprescu SN, Yue F, Qiu J, Brito LF, and Kuang S. Temporal Dynamics and Heterogeneity of Cell Populations during Skeletal Muscle Regeneration. iScience. 2020;23(4):100993.

57. Cosgrove BD, Gilbert PM, Porpiglia E, Mourkioti F, Lee SP, Corbel SY, et al. Rejuvenation of the muscle stem cell population restores strength to injured aged muscles. Nat Med. 2014;20(3):255–64.

58. Moyle LA, Cheng RY, Liu H, Davoudi S, Ferreira SA, Nissar AA, et al. Three-dimensional niche stiffness synergizes with Wnt7a to modulate the extent of satellite cell symmetric self-renewal divisions. Molecular Biology of the Cell. 2020;31(16):1703–13.

59. Chenette DM, Cadwallader AB, Antwine TL, Larkin LC, Wang J, Olwin BB, et al. Targeted mRNA Decay by RNA Binding Protein AUF1 Regulates Adult Muscle Stem Cell Fate, Promoting Skeletal Muscle Integrity. Cell Rep. 2016;16(5):1379–90.

60. Wheeler JR, Whitney ON, Vogler TO, Nguyen ED, Pawlikowski B, Lester E, et al. RNA-binding proteins direct myogenic cell fate decisions. Elife. 2022;11.

61. Maeda R, Kami D, Shikuma A, Suzuki Y, Taya T, Matoba S, et al. RNA decay in processing bodies is indispensable for adipogenesis. Cell Death Dis. 2021;12(4):285.

62. Li T, Shi Y, Wang P, Guachalla LM, Sun B, Joerss T, et al. Smg6/Est1 licenses embryonic stem cell differentiation via nonsense-mediated mRNA decay. Embo j. 2015;34(12):1630–47.

63. Abbadi D, Yang M, Chenette DM, Andrews JJ, and Schneider RJ. Muscle development and regeneration controlled by AUF1-mediated stage-specific degradation of fate-determining checkpoint mRNAs. Proceedings of the National Academy of Sciences. 2019;116(23):11285–90.

64. Blatt P, Wong-Deyrup SW, McCarthy A, Breznak S, Hurton MD, Upadhyay M, et al. RNA degradation is required for the germ-cell to maternal transition in Drosophila. Current Biology. 2021;31(14):2984–94.e7.

65. Solana J, Gamberi C, Mihaylova Y, Grosswendt S, Chen C, Lasko P, et al. The CCR4-NOT complex mediates deadenylation and degradation of stem cell mRNAs and promotes planarian stem cell differentiation. PLoS Genet. 2013;9(12):e1004003.

66. Lou CH, Chousal J, Goetz A, Shum EY, Brafman D, Liao X, et al. Nonsense-Mediated RNA Decay Influences Human Embryonic Stem Cell Fate. Stem Cell Reports. 2016;6(6):844–57.

67. Belair C, Sim S, Kim KY, Tanaka Y, Park IH, and Wolin SL. The RNA exosome nuclease complex regulates human embryonic stem cell differentiation. J Cell Biol. 2019;218(8):2564–82.

68. Di Stefano B, Luo EC, Haggerty C, Aigner S, Charlton J, Brumbaugh J, et al. The RNA Helicase DDX6 Controls Cellular Plasticity by Modulating P-Body Homeostasis. Cell Stem Cell. 2019;25(5):622–38.e13.

69. Hausburg MA, Doles JD, Clement SL, Cadwallader AB, Hall MN, Blackshear PJ, et al. Post-transcriptional regulation of satellite cell quiescence by TTP-mediated mRNA decay. Elife. 2015;4:e03390.

70. Chen Y-F, Li Y-SJ, Chou C-H, Chiew MY, Huang H-D, Ho JH-C, et al. Control of matrix stiffness promotes endodermal lineage specification by regulating SMAD2/3 via lncRNA LINC00458. Science Advances. 2020;6(6):eaay0264.

71. Brielle S, Bavli D, Motzik A, Kan-Tor Y, Sun X, Kozulin C, et al. Delineating the heterogeneity of matrix-directed differentiation toward soft and stiff tissue lineages via single-cell profiling. Proceedings of the National Academy of Sciences. 2021;118(19):e2016322118.

72. Trappmann B, Gautrot JE, Connelly JT, Strange DG, Li Y, Oyen ML, et al. Extracellular-matrix tethering regulates stem-cell fate. Nat Mater. 2012;11(7):642–9.

73. Selman M, and Pardo A. Fibroageing: An ageing pathological feature driven by dysregulated extracellular matrix-cell mechanobiology. Ageing Res Rev. 2021;70:101393.

74. Rivera T, Haggblom C, Cosconati S, and Karlseder J. A balance between elongation and trimming regulates telomere stability in stem cells. Nat Struct Mol Biol. 2017;24(1):30–9.

75. Liang J, Balachandra S, Ngo S, and O’Brien LE. Feedback regulation of steady-state epithelial turnover and organ size. Nature. 2017;548(7669):588-91.

76. Mendell JT, Sharifi NA, Meyers JL, Martinez-Murillo F, and Dietz HC. Nonsense surveillance regulates expression of diverse classes of mammalian transcripts and mutes genomic noise. Nat Genet. 2004;36(10):1073–8.

77. Goetz AE, and Wilkinson M. Stress and the nonsense-mediated RNA decay pathway. Cell Mol Life Sci. 2017;74(19):3509–31.

78. Pakos-Zebrucka K, Koryga I, Mnich K, Ljujic M, Samali A, and Gorman AM. The integrated stress response. EMBO Rep. 2016;17(10):1374–95.

79. Karam R, Lou CH, Kroeger H, Huang L, Lin JH, and Wilkinson MF. The unfolded protein response is shaped by the NMD pathway. EMBO Rep. 2015;16(5):599–609.

80. Deng M, Lin J, Nowsheen S, Liu T, Zhao Y, Villalta PW, et al. Extracellular matrix stiffness determines DNA repair efficiency and cellular sensitivity to genotoxic agents. Sci Adv. 2020;6(37).

81. Holstein EM, Clark KR, and Lydall D. Interplay between nonsense-mediated mRNA decay and DNA damage response pathways reveals that Stn1 and Ten1 are the key CST telomere-cap components. Cell Rep. 2014;7(4):1259–69.

82. Stoneley M, Harvey RF, Mulroney TE, Mordue R, Jukes-Jones R, Cain K, et al. Unresolved stalled ribosome complexes restrict cell-cycle progression after genotoxic stress. Molecular Cell. 2022;82(8):1557–72.e7.

83. Wong ES, Le Guezennec X, Demidov ON, Marshall NT, Wang ST, Krishnamurthy J, et al. p38MAPK controls expression of multiple cell cycle inhibitors and islet proliferation with advancing age. Dev Cell. 2009;17(1):142–9.

84. Perdiguero E, Ruiz-Bonilla V, Gresh L, Hui L, Ballestar E, Sousa-Victor P, et al. Genetic analysis of p38 MAP kinases in myogenesis: fundamental role of p38alpha in abrogating myoblast proliferation. Embo j. 2007;26(5):1245–56.

85. Wang C, Schmich F, Srivatsa S, Weidner J, Beerenwinkel N, and Spang A. Context-dependent deposition and regulation of mRNAs in P-bodies. eLife. 2018;7:e29815.

86. Pecori F, Kondo N, Ogura C, Miura T, Kume M, Minamijima Y, et al. Site-specific O-GlcNAcylation of Psme3 maintains mouse stem cell pluripotency by impairing P-body homeostasis. Cell Rep. 2021;36(2):109361.

87. Lipsitz LA, and Goldberger AL. Loss of ‘complexity’ and aging. Potential applications of fractals and chaos theory to senescence. Jama. 1992;267(13):1806–9.

88. Jaiswal S, and Ebert BL. Clonal hematopoiesis in human aging and disease. Science. 2019;366(6465).

89. Hernando-Herraez I, Evano B, Stubbs T, Commere P-H, Jan Bonder M, Clark S, et al. Ageing affects DNA methylation drift and transcriptional cell-to-cell variability in mouse muscle stem cells. Nature Communications. 2019;10(1):4361.

90. Clemens Z, Sivakumar S, Pius A, Sahu A, Shinde S, Mamiya H, et al. The biphasic and age-dependent impact of klotho on hallmarks of aging and skeletal muscle function. eLife. 2021;10:e61138.

91. Li X, Ploner A, Wang Y, Magnusson PKE, Reynolds C, Finkel D, et al. Longitudinal trajectories, correlations and mortality associations of nine biological ages across 20-years follow-up. eLife. 2020;9:e51507.

92. Tuck AC, Rankova A, Arpat AB, Liechti LA, Hess D, Iesmantavicius V, et al. Mammalian RNA Decay Pathways Are Highly Specialized and Widely Linked to Translation. Molecular Cell. 2020;77(6):1222–36.e13.

93. Smalec BM, Ietswaart R, Choquet K, McShane E, West ER, and Churchman LS. Genome-wide quantification of RNA flow across subcellular compartments reveals determinants of the mammalian transcript life cycle. bioRxiv. 2022:2022.08.21.504696.

94. Lashkevich KA, and Dmitriev SE. mRNA Targeting, Transport and Local Translation in Eukaryotic Cells: From the Classical View to a Diversity of New Concepts. Mol Biol. 2021;55(4):507–37.

95. Sahu A, Clemens ZJ, Shinde SN, Sivakumar S, Pius A, Bhatia A, et al. Regulation of aged skeletal muscle regeneration by circulating extracellular vesicles. Nature Aging. 2021;1(12):1148–61.

96. Stylianopoulos T, and Barocas VH. Multiscale, structure-based modeling for the elastic mechanical behavior of arterial walls. J Biomech Eng. 2007;129(4):611–8.

97. D’Amore A, Stella JA, Wagner WR, and Sacks MS. Characterization of the complete fiber network topology of planar fibrous tissues and scaffolds. Biomaterials. 2010;31(20):5345–54.

98. Zhao F, Xiong Y, Ito K, van Rietbergen B, and Hofmann S. Porous Geometry Guided Micro-mechanical Environment Within Scaffolds for Cell Mechanobiology Study in Bone Tissue Engineering. Front Bioeng Biotechnol. 2021;9:736489.

99. Smith LR, Hammers DW, Sweeney HL, and Barton ER. Increased collagen cross-linking is a signature of dystrophin-deficient muscle. Muscle Nerve. 2016;54(1):71–8.

100. Mula J, Lee JD, Liu F, Yang L, and Peterson CA. Automated image analysis of skeletal muscle fiber cross-sectional area. J Appl Physiol (1985). 2013;114(1):148-55.

101. Liu L, Cheung TH, Charville GW, and Rando TA. Isolation of skeletal muscle stem cells by fluorescence-activated cell sorting. Nat Protoc. 2015;10(10):1612–24.

102. Soneson C, and Robinson MD. Bias, robustness and scalability in single-cell differential expression analysis. Nat Methods. 2018;15(4):255–61.

103. Hettinger ZR, Wen Y, Peck BD, Hamagata K, Confides AL, Van Pelt DW, et al. Mechanotherapy Reprograms Aged Muscle Stromal Cells to Remodel the Extracellular Matrix during Recovery from Disuse. Function (Oxf). 2022;3(3):zqac015.

104. Rocheteau P, Gayraud-Morel B, Siegl-Cachedenier I, Blasco MA, and Tajbakhsh S. A subpopulation of adult skeletal muscle stem cells retains all template DNA strands after cell division. Cell. 2012;148(1-2):112–25.

105. Xing J. Reconstructing data-driven governing equations for cell phenotypic transitions: integration of data science and systems biology. Physical Biology. 2022;19(6):061001.

106. Reimand J, Isserlin R, Voisin V, Kucera M, Tannus-Lopes C, Rostamianfar A, et al. Pathway enrichment analysis and visualization of omics data using g:Profiler, GSEA, Cytoscape and EnrichmentMap. Nat Protoc. 2019;14(2):482–517.

107. Svensson V, Natarajan KN, Ly L-H, Miragaia RJ, Labalette C, Macaulay IC, et al. Power analysis of single-cell RNA-sequencing experiments. Nature Methods. 2017;14(4):381–7.

